# DksA-dependent regulation of RpoS contributes to *Borrelia burgdorferi* tick-borne transmission and mammalian infectivity

**DOI:** 10.1101/2020.11.04.367946

**Authors:** William K. Boyle, Crystal L. Richards, Daniel P. Dulebohn, Amanda K. Zalud, Jeff A. Shaw, Sándor Lovas, Frank C. Gherardini, Travis J. Bourret

**Affiliations:** Department of Medical Microbiology and Immunology, Creighton University, Omaha, NE 68105; Laboratory of Bacteriology, Gene Regulation Section, Division of Intramural Research, Rocky Mountain Laboratories, National Institute of Allergy and Infectious Diseases, National Institutes of Health, Hamilton, MT 59840; Department of Biomedical Sciences, Creighton University, Omaha, NE 68105

## Abstract

Throughout its enzootic cycle, the Lyme disease spirochete *Borreliella (Borrelia) burgdorferi*, senses and responds to changes in its environment by using a small repertoire of transcription factors which coordinate the expression of genes required for infection of *Ixodes* ticks and various mammalian hosts. Among these transcription factors, the DnaK suppressor protein (DksA) plays a pivotal role in regulating gene expression in *B. burgdorferi* during periods of nutrient limitation and is required for mammalian infectivity. In many pathogenic bacteria, the gene regulatory activity of DksA along with the alarmone guanosine penta- and tetra-phosphate ((p)ppGpp) coordinates the stringent response to various environmental stresses including nutrient limitation. In this study, we sought to characterize the role of DksA in regulating the transcriptional activity of RNA polymerase and in the regulation of RpoS-dependent gene expression required for *B. burgdorferi* infectivity. Using *in vitro* transcription assays, we observed recombinant DksA inhibits RpoD-dependent transcription by *B. burgdorferi* RNA polymerase independent of ppGpp Additionally, we determined the pH-inducible expression of RpoS-dependent genes relies on DksA, but is independent of (p)ppGpp produced by Rel_bbu_. Subsequent transcriptomic and western blot assays indicated DksA regulates the expression of BBD18, a protein previously implicated in the post-transcriptional regulation of RpoS. Moreover, we observed DksA was required for infection of mice following intraperitoneal inoculation or for transmission of *B. burgdorferi* by *Ixodes scapularis* nymphs. Together, these data suggest DksA plays a central role in coordinating transcriptional responses of *B. burgdorferi* required for infectivity through its interactions with RNA polymerase and post-transcriptional control of RpoS.

**Author Summary:** Lyme disease, caused by the spirochetal bacteria *Borrelia burgdorferi*, is the most common vector-borne illness in North America. The ability of *B. burgdorferi* to establish infection is predicated by its ability to coordinate the expression of virulence factors in response to diverse environmental stimuli encountered within *Ixodes* ticks and mammalian hosts. Previous studies have shown an essential role for the alternative sigma factor RpoS in regulating the expression of genes required for the successful transmission of *B. burgdorferi* by *Ixodes* ticks and infection of mammalian hosts. The DnaK suppressor protein (DksA) is a global gene regulator in *B. burgdorferi* that also contributes to the expression of RpoS-dependent genes. In this study, we determined DksA exerts its gene regulatory function through direct interactions with the *B. burgdorferi* RNA polymerase using *in vitro* transcription assays and controls the expression of RpoS-dependent genes required for mammalian infection by post-transcriptionally regulating cellular levels of RpoS. Our results demonstrate the utility of *in vitro* transcription assays to determine how gene regulatory proteins like DksA control gene expression in *B. burgdorferi*, and reveal a novel role for DksA in the infectious cycle of *B. burgdorferi*.

## INTRODUCTION

*Borrelia burgdorferi*, a causative agent of Lyme disease, is the most predominant vector-borne pathogen in the United States (1, 2). *B. burgdorferi* is an extracellular pathogen that traverses diverse environments within its tick vector, *Ixodes scapularis*, and mammalian hosts. During blood meal acquisition, *B. burgdorferi* residing within *I. scapularis* midguts are subjected to changes in temperature, osmolarity, pH, and host-derived reactive oxygen and nitrogen species (3-5). In addition to these potential environmental stressors, the midgut concentrates components of mammalian blood, including nutrients, organic acids, host derived-antibodies and immune complement components during and following tick-feeding (6-8). *B. burgdorferi* adapts to this dynamic environment within feeding ticks by shifting gene expression and replicating rapidly prior to transiting from the tick midgut to mammalian hosts.

The DnaK suppressor protein (DksA) and the metabolite guanosine penta- and tetra phosphate ((p)ppGpp) produced by Rel_bbu_ coordinate the stringent response of *B. burgdorferi* by regulating the transcription of genes in response to nutrient limitation (9-11). Furthermore, there is an extensive overlap in genes under the control of DksA and (p)ppGpp in *B. burgdorferi* (11). DksA and (p)ppGpp both control genes involved in protein synthesis, nutrient uptake pathways, as well as several lipoproteins. Additionally, DksA and (p)ppGpp control the expression of genes encoding products that facilitate glycerol and chitobiose utilization, along with genes encoding oligopeptide transporters, which are essential for the adaptation of *B. burgdorferi* during infection of *I. scapularis* (12-14). These observations suggest DksA and (p)ppGpp work synergistically to regulate transcription by RNA polymerase in *B. burgdorferi* (11) and are consistent with the generation of a stringent response due to the overlapping regulatory roles of DksA and (p)ppGpp in other organisms (15-18) where these regulators control the response to stationary phase growth and nutrient limiting conditions (19, 20). DksA derived from other bacteria such as *Escherichia coli* interacts with (p)ppGpp *in vitro* which can fundamentally change the direction and magnitude of gene regulation from specific gene promoters in *E. coli* using *in vitro* transcription reactions (21-24).

The nature of the stringent response in *B. burgdorferi* is not fully understood. *B. burgdorferi* requires a complex, nutrient-rich medium for growth and what constitutes nutrient limitation in *B. burgdorferi* is not well-defined. Previous attempts to study the stringent response were carried out by altering multiple environmental and metabolic factors simultaneously, making it difficult to interpret which key factors and metabolites played a role in the generation of a stringent response by *B. burgdorferi* (9, 11, 25-27). While limitation of nutrients including fatty acids, carbohydrates, or amino acids can activate the stringent response in bacteria, responses involving DksA and (p)ppGpp are also triggered by environmental changes, such as low pH and the presence of reactive nitrogen species (28-30). For example, in *E. coli*, the adaptation to an acidic extracellular environment is aided by the regulation of the alternative sigma factor RpoS by DksA and DksA appears to be a key component of the response of *E. coli* to acid stress (31, 32). We recently observed *dksA*-deficient *B. burgdorferi* strains showed reduced expression of genes under the control of the alternative sigma factor RpoS (11). RpoS plays a gatekeeper function in the infectious cycle of *B. burgdorferi* by coordinating the expression of a substantial portion of genes during the transmission of *B. burgdorferi* from ticks to mammalian hosts.

During *in vitro* growth, RpoS is robustly expressed by *B. burgdorferi* in Barbour-Stoenner-Kelly II (BSKII) growth media under mildly acidic conditions designed to mimic the pH and organic acid stresses encountered within the *I. scapularis* midgut (5, 33, 34). The *I. scapularis* midgut lumen is slightly acidic (pH 6.8) and contains millimolar concentrations of organic acids following blood meal acquisition by ticks from mammalian hosts (5, 35). Although the mechanisms underlying the activation of the stringent response in *B. burgdorferi* remain unclear, the role DksA plays in the regulation of RpoS may yield answers to what environmental conditions are important for unlocking DksA-dependent gene regulation.

RpoS regulates the expression of genes encoding lipoproteins, which contribute to mammalian infection, such as *bba66*, decorin binding protein (*dbpA*), and outer surface protein C (*ospC*) (36-38). Each of these genes are highly expressed by *B. burgdorferi* residing within the *I. scapularis* midgut during transmission, and RpoS-dependent expression of these genes is required by *B. burgdorferi* to adapt and transition to the mammalian host environment (39-41). RpoS itself is subject to both transcriptional and post-transcriptional regulation in *B. burgdorferi* (42, 43). The alternative sigma factor RpoN, drives the expression of a short *rpoS* transcript, which leads to the accumulation of RpoS (44, 45). The RpoN-dependent expression of *rpoS* relies on additional transcription factors, including the response regulator Rrp2 and the ferric uptake regulator homolog, BosR. Additionally, *rpoS* expression is inhibited by the *Borrelia* adaptation protein (BadR) (45-52). Alternatively, accumulation of RpoS can be driven by the expression of a long transcript through a RpoN-independent mechanism (53). Post-transcriptional control of *rpoS* is regulated by small RNA-dependent mechanisms and by BBD18, a protein that contributes to RpoS degradation (54-57). However, the mechanisms underlying the ability of *B. burgdorferi* to sense shifts in pH or organic acid stress to regulate *rpoS*/RpoS remain unclear.

Further characterization of DksA-dependent gene regulation is required to reconcile the nature of the stringent response and the possible role of DksA in controlling RpoS. In this study, we used both biochemical and genetic approaches to dissect the gene regulatory role played by DksA and its relationship with (p)ppGpp in *B. burgdorferi*. In our genetic approach, we assessed pH- and acetate-dependent responses of *B. burgdorferi* in the presence and absence of DksA and the (p)ppGpp synthase (Rel_bbu_) because previous studies suggested DksA plays a much larger role in regulating RpoS (11, 25). To test the hypothesis that a synergy exists between DksA (p)ppGpp in regulating gene transcription, we used a newly developed *in vitro* transcription assay system using the *B. burgdorferi* RNA polymerase (58). Due to the important regulatory role of DksA in controlling factors required for the transmission of *B. burgdorferi*, we determined the contribution of DksA to tick-borne transmission and mammalian infectivity. We show that DksA is required for murine infection following intraperitoneal inoculation as well as transmission by artificially-infected *I. scapularis* nymphs to mice. Additionally, we found DksA and (p)ppGpp play independent roles in controlling RpoS levels in *B. burgdorferi*. DksA contributes to the post-transcriptional regulation of RpoS, possibly through its control of BBD18, and the pH-dependent expression of RpoS-regulated genes. We observed (p)ppGpp is produced at high levels in response to growth of *B. burgdorferi* under acidic conditions but is not required for RpoS expression. Collectively, the results of this study provide evidence that weak acidic environments activate the *B. burgdorferi* stringent response and that DksA is an indispensable transcriptional regulator coordinating the expression of genes required for mammalian infection.

## RESULTS

### DksA acts as a transcriptional repressor

Previous work by our laboratory characterized the role of DksA in the *B. burgdorferi* stringent response (11) and found DksA plays an important global regulatory role that partially overlapped with (p)ppGpp-dependent gene regulation in nutrient limited conditions. To further dissect the role of DksA, ppGpp, the interaction of these two regulatory factors and define the molecular basis of transcriptional regulation enacted by these regulators we analyzed structural and functional aspects of DksA in vitro. Similar to other DksA homologs, *B. burgdorferi* DksA is predicted to be an α-helix rich peptide with a carboxy-terminal zinc-finger motif (11). To better understand the molecular characteristics of the *B. burgdorferi* DksA, we carried out experiments to determine alpha helicity and zinc binding using recombinant *B. burgdorferi* DksA protein. DksA was purified to apparent homogeneity using a maltose affinity purification system (Fig. 1A), and the α-helicity of the protein was measured by circular dichroism (Fig. 1B). Recombinant DksA had a high positive molar ellipticity maximum at 195 nm and negative molar ellipticity maxima at 208 and 222 nm wavelengths, indicating *B. burgdorferi* DksA is an α-helix rich protein (Fig. 1C). The putative zinc finger domain of DksA is comprised of four cysteinyl residues (C91, C94, C112, and C115), suggesting *B. burgdorferi* DksA falls within a redox-active group of DksA proteins (59, 60). The release of Zn^2+^ from DksA following exposure to increasing concentrations of H_2_O_2_ was tested using the Zn^2+^-sensitive dye 4-(2-pyridylazo)resorcinol (PAR) (Fig. 1D). Incubation of 100 μmol DksA with 1.25 mM H_2_O_2_ for 1.5 hours resulted in the release of 40 μmol Zn^2+^, presumably due to the release of Zn^2+^ following the oxidation of the Cys residues comprising the zinc finger. Together, these results indicate recombinant *B. burgdorferi* DksA is an α-helix rich, Zn^2+^-containing polypeptide, similar to other characterized DksA proteins.

**Figure 1.**
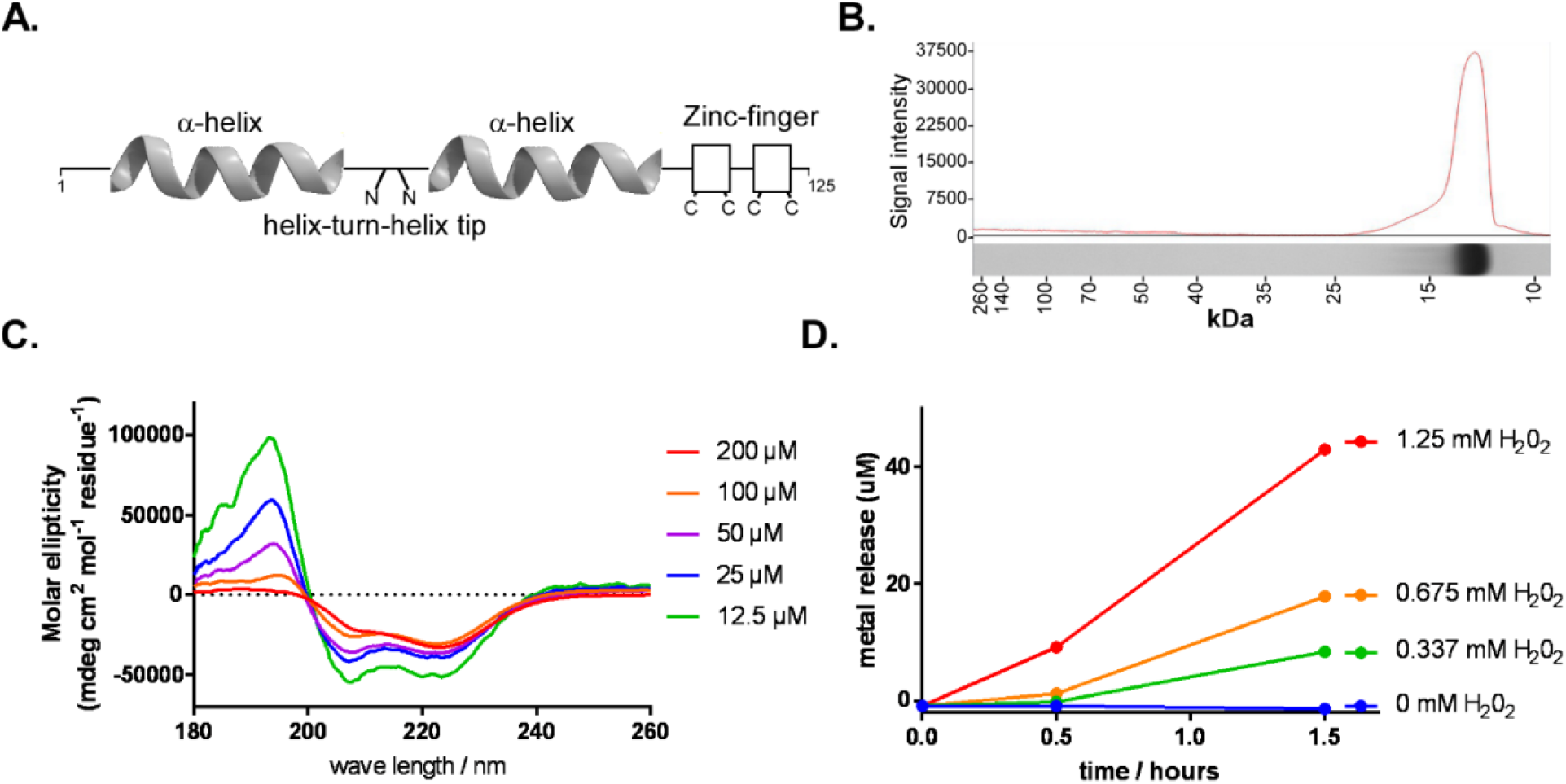
Measurement of *B. burgdorferi* DksA α-helicity and Zn^2+^-binding. (A) Schematic of DksA predicted structure. (B) DksA purified to apparent homogeneity. DksA purity was assessed by Coomassie stained SDS-PAGE gel. (C) Circular dichroism spectra of recombinant *B. burgdorferi* DksA measured in 15 mM sodium phosphate buffer pH 7.4 and 100 mM NaCl. (D) Zn^2+^ release from 100 uM of DksA mixed with various concentrations of H202 was measured by a zinc sensitive dye 4-(2-pyridylazo)resorcinol (PAR). Increase in Zn^2+^ release with higher levels of H202 suggest 4-Cys zinc finger is capable of coordinating Zn^2+^.

Two strong RpoD-dependent promoters (*flgBp* and *rplUp*) and five promoters regulated by the stringent response (11, 25) (*clpCp, glpFp, groLp, napAp*, and *nagAp*) were selected to determine the impact of recombinant *B. burgdorferi* DksA or ppGpp on RNA polymerase activity from these promoters. Varying levels of ppGpp (0, 20, and 200 μM) and DksA (0, 50, and 500 nM) were added to *in vitro* transcription reactions to test for their effects on transcriptional initiation by RNA polymerase as shown in previous investigations of other *in vitro* transcription systems (29, 61, 62). *In vitro* transcription reactions with RNA polymerase supplemented with recombinant RpoD along with varying concentrations of DksA and/or ppGpp were prepared in parallel. Reactions were then initiated by the addition of double-stranded linear DNA templates encoding the promoters of interest and the reaction proceeded for five minutes at 37 °C. Following termination, RNA products were separated by gel electrophoresis and the relative incorporation of α-^32^P-ATP was detected by phosphor imaging (Fig. 2A). The addition of 500 nM of DksA (roughly 24 molecules DksA: 1 molecule RNA polymerase) led to a significant reduction of RNA polymerase activity from all of the tested promoters (Fig. 2B). In contrast, the addition of ppGpp did not lead to significant changes in RNA polymerase activity from any of the promoters tested (Fig 2C), indicating ppGpp had no direct effects on *B. burgdorferi* RNA polymerase activity *in vitro*.

**Figure 2.**
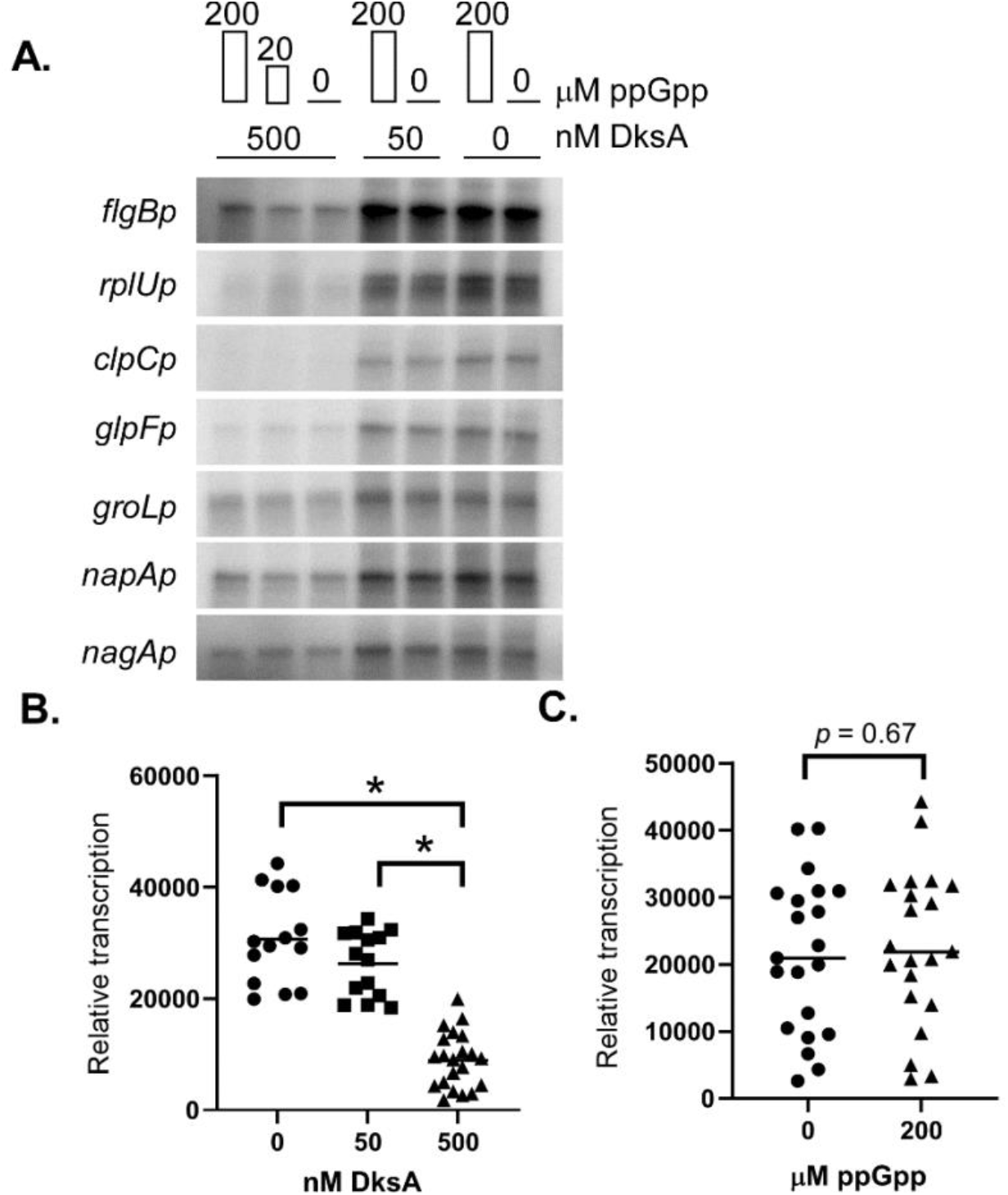
*In vitro* transcription reactions to assay ppGpp and DksA effects on transcription initiation from seven *B. burgdorferi* promoters. *In vitro* transcription reactions initiated on seven RpoD-dependent promoters were prepared containing 500, 50, or 0 nM DksA and 200, 20, or 0 μM of ppGpp. (A) RNA was separated by gel electrophoresis on a 10% urea gel are shown. First, fourth, and sixth lanes contained 200 μM of ppGpp. Second and fifth lane contained 20 μM ppGpp. Phosphor screen signals from all *in vitro* transcription were pooled to evaluate (B) DksA-dependent effects on transcription using a non-parametric ranked Kruskal–Wallis multiple comparisons test and (C) ppGpp-dependent effects using non-parametric ranked Mann-Whitney comparison test. Asterisk indicates p-value < 0.001 in multiple comparison testing.

The promoters used in this study displayed varying levels of transcription in the absence of DksA, as illustrated by the higher levels of transcription from the *flgBp* compared to *napAp*. However, transcription from *flgBp* and *napAp* appeared similar in the presence of 500 nM DksA. To test the promoter-dependent effects of DksA on transcription, a broader range of DksA concentrations were tested for *in vitro* transcription reactions initiated on *flgBp* or *napAp* (Fig. 3). DksA affected both promoters in a concentration-dependent manner, demonstrated by reduced transcription from each promoter with the addition of increasing concentrations of DksA in the reaction. The addition of only 250 nM DksA (roughly 12 molecules DksA: 1 molecule RNA polymerase) resulted in a 2 – 4 fold reduction of RNA polymerase activity from both promoters (Fig. 3A). Together, these data support a role for DksA as a transcriptional regulator that directly affects RNA polymerase activity from various *B. burgdorferi* promoters.

**Figure 3.**
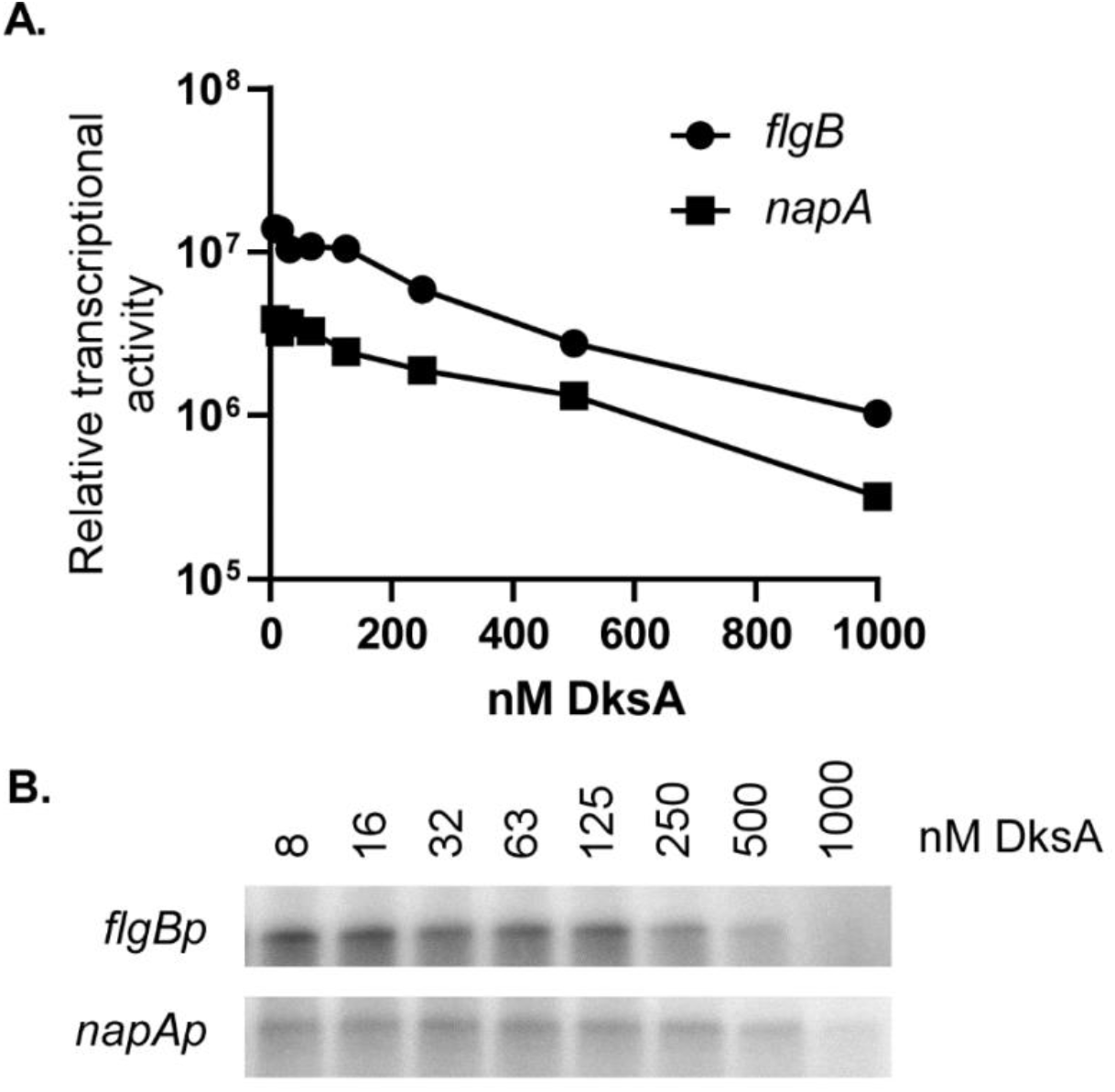
Test of DksA concentration-dependent regulation of *flgB* and *napA*. Relative RNA product amounts after *in vitro* transcription reactions containing various concentrations (8 – 1000 nM) of DksA. Two promoters *flgBp* and *napAp* which appeared to have different levels of response to DksA in Figure 1 were tested. Signal intensity as measured by isotopically labeled RNA and determined by densitometry (A) of phosphor screen images (B) are shown.

### DksA contributes to post-transcriptional regulation of RpoS

Previous studies have demonstrated *B. burgdorferi* strains with mutations of *dksA* show a low levels of expression of genes under the control of the Rrp2-RpoN-RpoS regulatory cascade (e.g. *dbpA, ospC, bba66*) compared to wild-type controls (11, 63). The expression of genes under the control of the Rrp2-RpoN-RpoS pathway are known to increase during growth in mildly acidic BSKII, pH 6.7 medium and with the addition of organic acids such as acetate (5, 33, 64). To determine whether DksA contributes to the pH-dependent expression of RpoS, western blot analyses were carried out using whole cell lysates collected from mid-log phase cultures of wild-type, Δ*rel*_bbu_, and Δ*dksA B. burgdorferi* strains, along with a chromosomally complemented strain (Δ*dksA* cDksA), grown in either BSKII, pH 7.6 or BSKII, pH 6.7 + 20 mM acetate. Consistent with previous studies (49, 65), the expression of RpoS was higher in cultures of wild-type *B. burgdorferi* grown in pH 6.7 + 20 mM acetate compared to cultures grown in BSKII, pH 7.6 (Fig. 4A). In contrast, RpoS was undetectable in the Δ*dksA* strain, while patterns of RpoS expression in the Δ*rel*_bbu_ and Δ*dksA* cDksA strains were comparable to wild-type controls. Similarly, the expression of OspC was increased in all but the Δ*dksA* strain in response to growth in pH 6.7 + 20 mM acetate (Fig. 4B).

**Figure 4.**
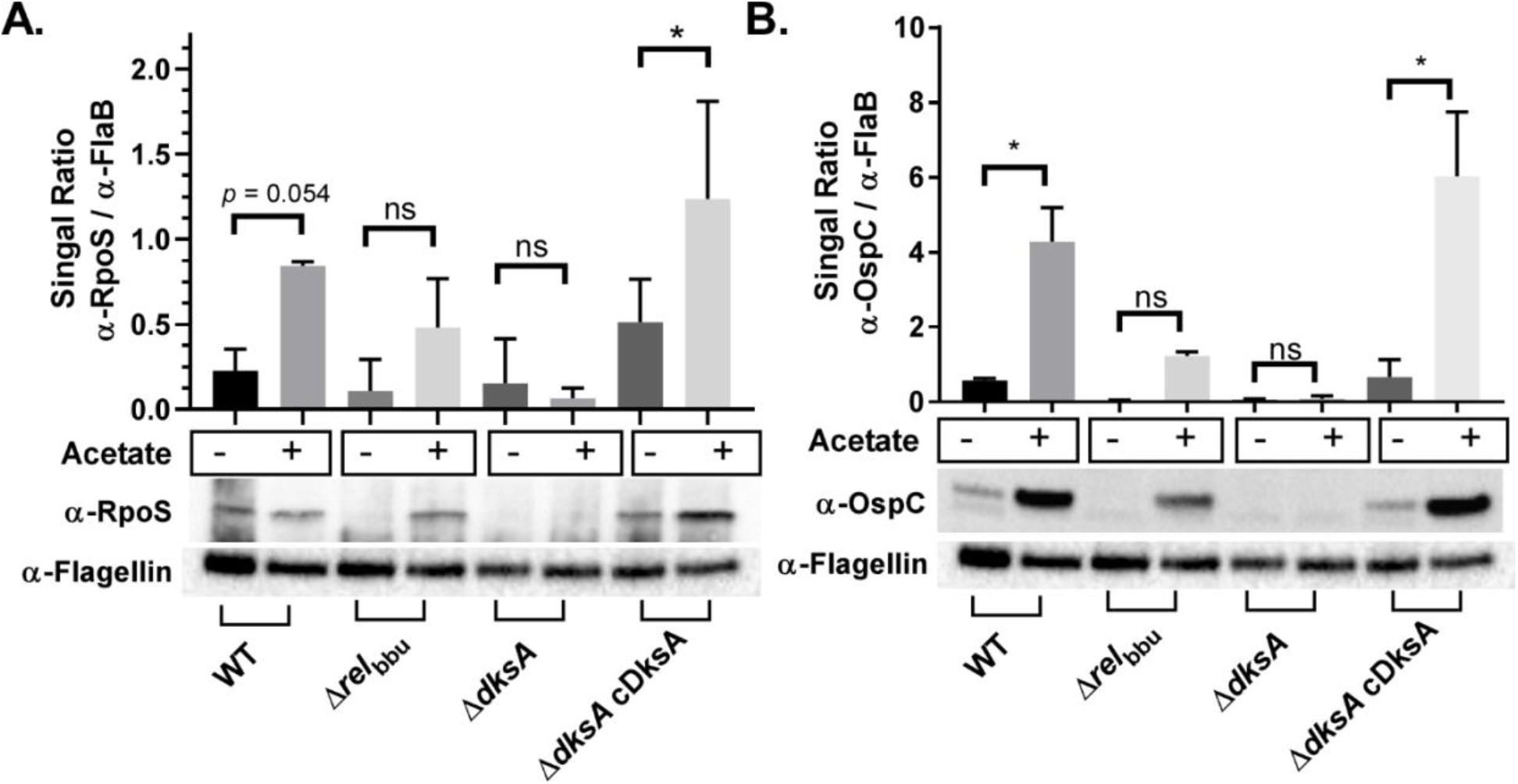
Relative expression of RpoS and OspC in wild-type (WT), Δ*relbbu*, Δ*dksA*, and Δ*dksA* cDksA (cDksA) *B. burgdorferi* strains at mid-logarithmic growth. Whole cell lysates were collected from *B. burgdorferi* 297 strain cultured to 5 × 10^7^ spirochetes/ mL in either BSKII-pH 7.6 or BSKII-pH 6.7 with 20 mM acetate. Expression of RpoS and OspC protein levels were determined by normalizing densitometry of chemiluminescence signals obtained from western blot to signals obtained from flagellin. Data are representative of 3 replicate experiments. Asterisk indicate adjusted *p*-values < 0.05 in a Sidak multiple comparisons test.

Next, we investigated whether DksA contributed to the transcription of *rpoS* and the RpoS-regulated gene *ospC* in response to growth in mildly acidic conditions. All three strains, wild-type, Δ*rel*_bbu_, and Δ*dksA B. burgdorferi*, showed increased expression of *rpoS* when grown in BSKII, pH 6.7 + 20 mM acetate compared to cultures grown in BSKII, pH 7.6 (Fig. 5A), suggesting DksA is not required for the pH-dependent increases in *rpoS* transcription. We observed pH-dependent increases in *ospC* in both wild-type and Δ*rel*_bbu_ strains, while the Δ*dksA* strain was unable to induce the expression of *ospC* in response to growth under acidic conditions, consistent with the patterns of OspC protein levels observed in each strain. The observation that pH-induced expression of *rpoS* remains intact in the Δ*dksA* strain was further supported by results from experiments where RNA was collected from mid-logarithmic and stationary phase cultures to detect the increase of *rpoS* expression at stationary phase (supplemental Fig. 1). At stationary phase, wild-type and Δ*dksA* cultures showed increased *rpoS* expression, while the Δ*dksA* strain did not express RpoS-regulated genes *dbpA, ospC*, and *bba66* at levels observed in wild-type controls as determined by a multiple comparisons test (*p-*value < 0.05). These results support a role for DksA in the post-transcriptional regulation of *rpoS*/RpoS that is independent of Rel_bbu_.

**Figure 5.**
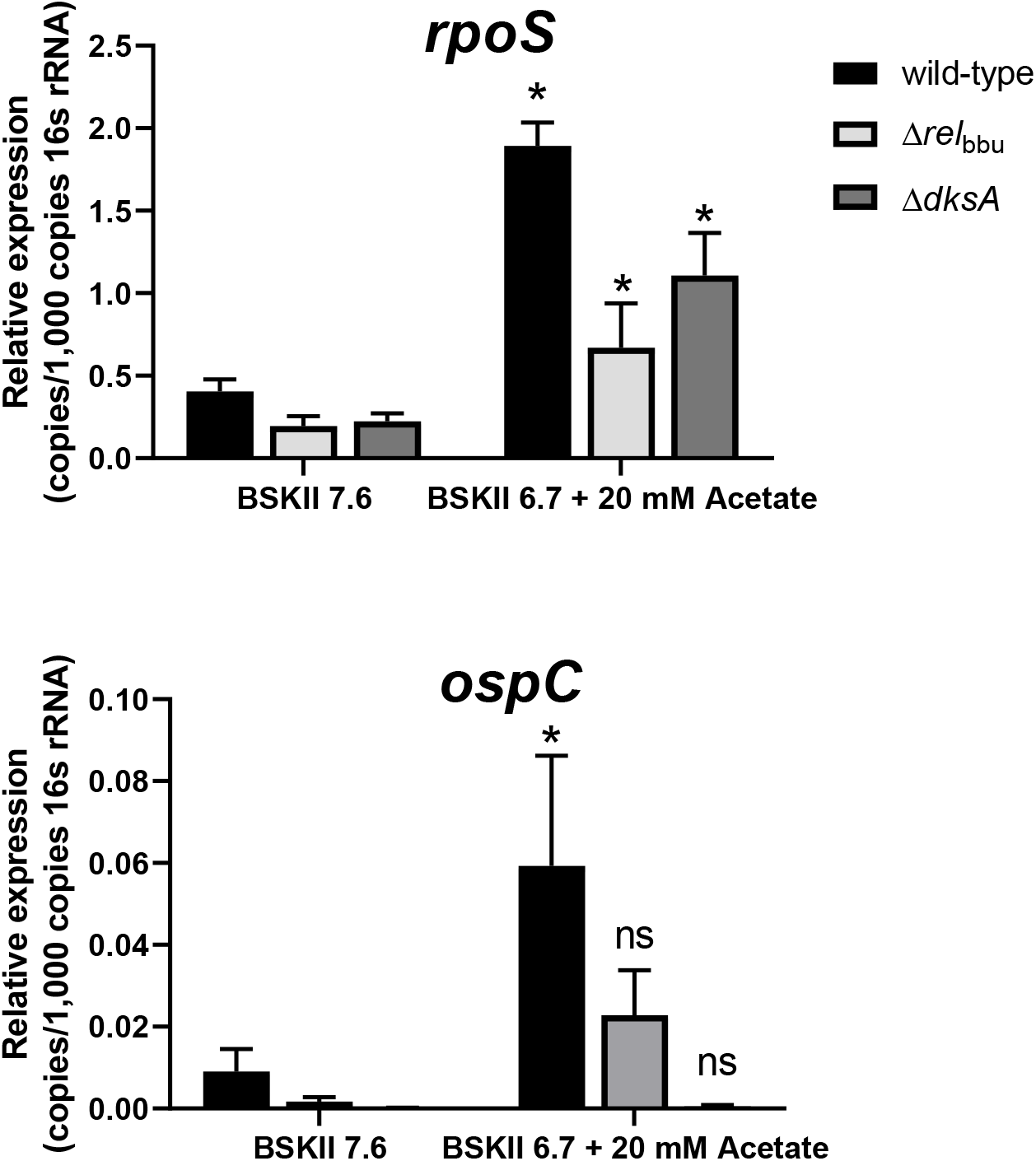
RT-qPCR analysis of *rpoS* and *ospC* transcripts in wild-type, Δ*rel*_bbu_ and Δ*dksA* strains. RT-qPCR was performed on RNA extracted from wild-type (WT), Δ *rel*_bbu_, and Δ*dksA* mid-logarithmic phase cultures in BSKII 7.6 or BSKII 6.7 + 20 mM acetate. Relative expression of *ospC* and *rpoS* were normalized to 16s rRNA. Error bars represent standard deviation calculated from four biological replicates. Asterisk indicates *p*-value < 0.05 for relative expression between BSKII 7.6 and BSKII 6.7 + 20 mM acetate conditions in a Holm-Sidak multiple comparisons test.

In a recent study by Mason, et al., *B. burgdorferi* DksA was implicated in the post-transcriptional regulation of *rpoS*, possibly through the transcriptional regulation of the sRNA chaperone encoded by *hfq* and along with the protease *clpX/clpP/clpP2*. In this study we assayed another known post-transcriptional regulator of RpoS in *B. burgdorferi*, BBD18. Increased levels of BBD18 are correlated with reduced levels of RpoS (56). Our previous transcriptomic study indicated increased expression of *bbd18* in the Δ*dksA* strain (11). We quantified the relative levels of BBD18 by western blot to determine if elevated levels of BBD18 correspond with a reduction in RpoS. BBD18 expression levels were lower in response to growth in BSK II pH 6.7 + 20 mM acetate in all strains except the Δ*dksA* strain (Fig. 6). Results indicate the Δ*dksA* strain is unable to downregulate the expression of BBD18 in response to mildly acidic growth conditions, which corresponds with the reduced levels of RpoS shown in Fig. 1. Additionally, the alteration of the growth rate of *B. burgdorferi* has been proposed as an alternative mechanism for regulating cellular levels of OspC based on protein accumulation (66). To determine if growth rates are significantly altered in the *B. burgdorferi* Δ*dksA*- and Δ*rel*_bbu_ mutant strains, BSKII containing 0, 30, and 100 mM acetate, at pH 7.6, 6.1, and 4.6 respectively, were inoculated with 10^6^ spirochetes / ml and monitored for cell density over time (Fig. 7). Similar growth rates were observed for each strain in BSKII + 30 mM acetate, indicating differences in OspC expression could not be explained by differences in the growth of these strains. These results suggest DksA controls BBD18-dependent post-transcriptional regulation of *rpoS*/RpoS, independent of the growth rate of *B. burgdorferi*.

**Figure 6.**
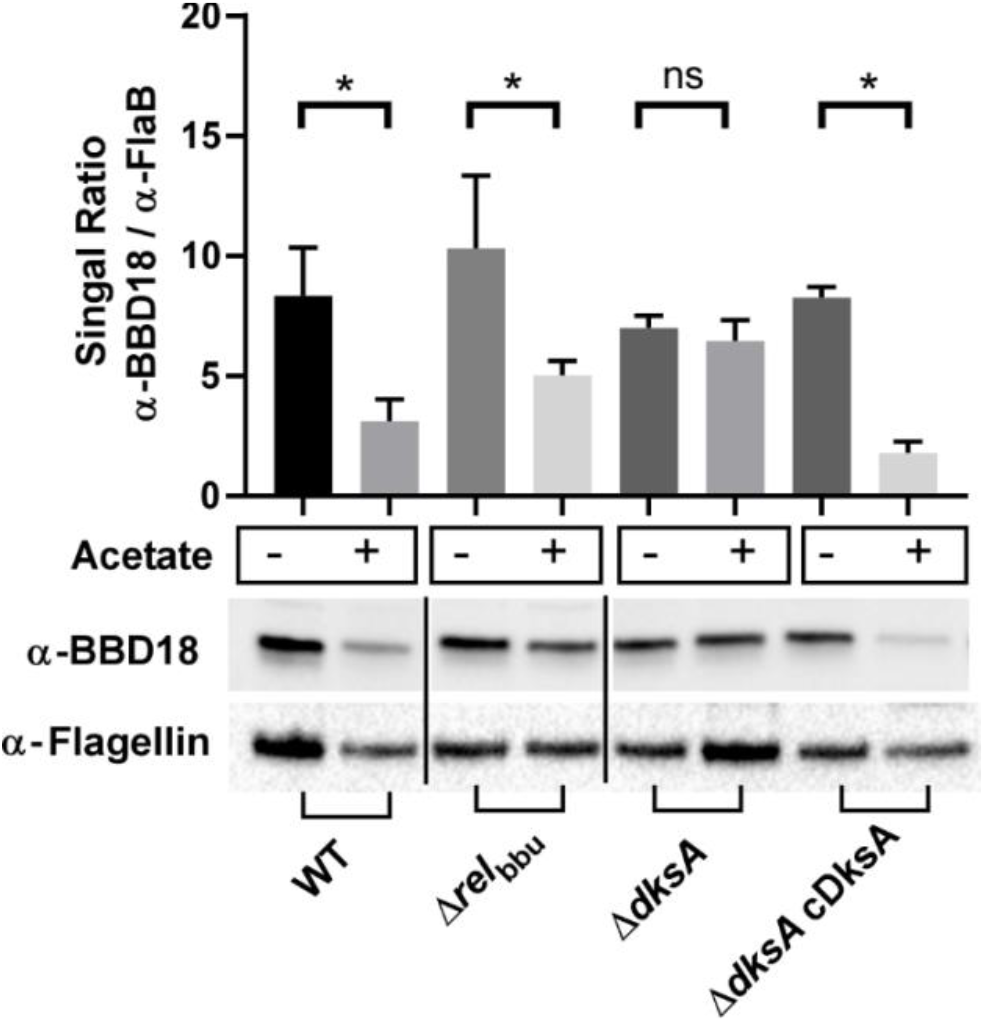
Relative expression of BBD18 by *B. burgdorferi* in response to growth in BSKII pH 7.6 and BSKII pH 6.7 + 20mM acetate. Whole cell lysates were collected from 297 wild-type (WT), Δ*rel*_bbu_, Δ*dksA*, and chromosomally complemented Δ*dksA (ΔdksA* cDksA) *B. burgdorferi* strains cultured to 5 × 10^7^ spirochetes / mL density in either BSKII-pH 7.6 or BSKII-pH 6.7 with 20 mM acetate. Expression of BBD18 protein levels were determined by densitometry of chemiluminescence signals and signals were normalized to expression of FlaB. Data are representative of 3 replicate experiments. Asterisk indicate adjusted *p*-values < 0.05 in a Sidak multiple comparisons test.

**Figure 7.**
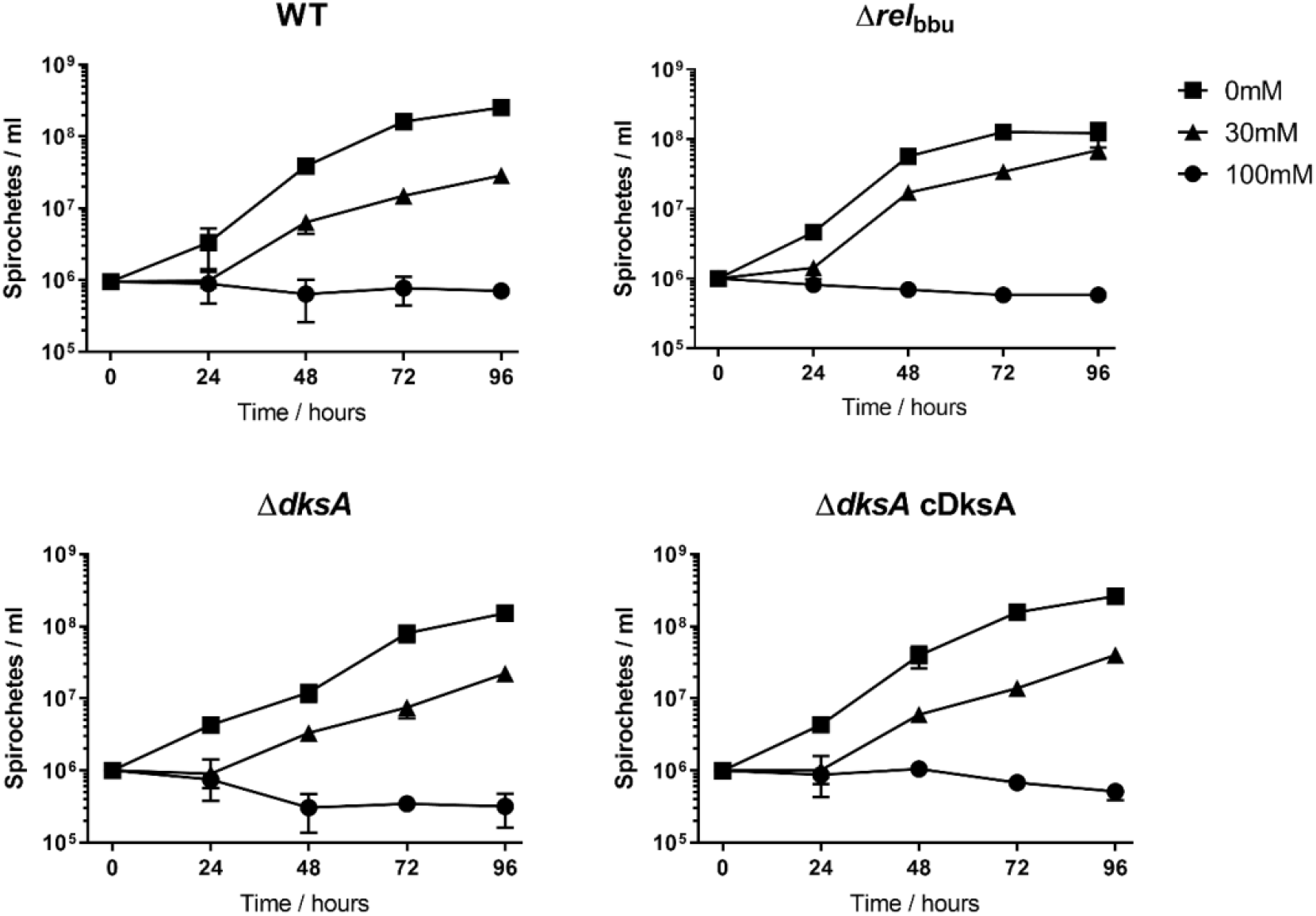
Growth rates of *B. burgdorferi* in acidic environment. *B. burgdorferi* 297 wild-type (WT), Δ*rel*_bbu_, Δ*dksA*, and chromosomally complemented Δ*dksA* strain (Δ*dksA* cDksA) were cultured in BSK II containing 0 (square), 30 (triangle), and 100 mM (circle) of acetate. Media pH was not compensated for the addition of acetic acid and 0, 30, and 100 mM medium also represent growth in pH 7.6, 6.1, and 4.6 respectively. Growth curves are representative of 3 replicate cultures and error bars represent standard deviation.

### DksA is required for mammalian infection

A recent study by Mason, et. al. performed concurrently with our own investigation showed DksA is required for the infection of mice following intradermal inoculation with *B. burgdorferi* (63). Consistent with the data shown in our study, the requirement of DksA for mammalian infection was linked with dysregulation of the Rrp2-RpoN-RpoS pathway. Therefore, we independently assessed the role of DksA in mammalian infection by intraperitoneal inoculation of naïve RML mice with 5 × 10^7^ *B. burgdorferi* 297 wild-type, Δ*dksA*, or Δ*dksA* cDksA *B. burgdorferi* strains. At 23 days post inoculation, the ear, joint, and bladder tissues were collected from mice and transferred to BSKII medium to detect the presence of *Borrelia*. Viable spirochetes were isolated from all cultured tissues of the mice injected with wild-type and Δ*dksA* cDksA strains; however, none of the tissues from mice inoculated with the Δ*dksA* strain showed spirochete outgrowth (Table 1). Additionally, seroconversion of mice infected with the *B. burgdorferi* strains was tested against wild-type *B. burgdorferi* cell lysates by western blot using serum collected from mice 23 days post infection. Mice infected with wild-type and Δ*dksA* cDksA strains showed seroconversion, while mice infected with the Δ*dksA* strain did not (Supplemental Fig. 2). Collectively, these data indicate DksA is required for *B. burgdorferi* to establish infection in mice.

**Table 1.** Evaluation of the potential for DksA-deficient *B. burgdorferi* to cause infection.

| Stain | Ear | Joint | Bladder | Control<sup>(a)</sup> | Serology |
| --- | --- | --- | --- | --- | --- |
| Wild-type | 4/4<sup>(b)</sup> | 5/5 | 5/5 | 1/1 | 5/5 |
| $\Delta dksA$ | 0/5 | 0/5 | 0/5 | 1/1 | 0/5 |
| $\Delta dksA$ cDksA | 5/5 | 5/5 | 5/5 | 1/1 | 5/5 |

Numbers indicate ratio of *B. burgdorferi* positive cultures from various tissue sites, ear, joint, or heart and positive serology over the number of individual mice tested. (a) control tubes tested the ability of each *B. burgdorferi* strain to grow in media containing antibiotics mixture. (b) one culture was lost due to contamination.

Our previous transcriptomic study showed DksA controls the expression of a wide variety of genes that have previously been shown to contribute to colonization and transmission by *I. scapularis*. To more robustly test the role of DksA in host and vector infection and colonization, we artificially infected *I. scapularis* larvae by submersion in wild-type, Δ*dksA*, or Δ*dksA* cDksA cultures grown to a density of 5 × 10^7^ spirochetes · ml^-1^ in BSKII and tested the ability of the larvae to transmit the three strains to naïve mice. Approximately 200 artificially infected larval stage ticks with either the wild-type, Δ*dksA*, or Δ*dksA* cDksA strains were allowed to feed on mice. Spirochetes in larval ticks were quantified, 3 – 5 days following the larval bloodmeal, by plating a subset of homogenized ticks on semi solid BSK media. An additional subset of ticks were analyzed following the molt of larvae to nymphs roughly 6 weeks after feeding (Supplemental Fig. 3). *I. scapularis* larvae and nymphs colonized by the Δ*dksA* or Δ*dksA* cDksA strains showed lower spirochete numbers compared to wild-type controls, however, statistical analysis by one-way ANOVA and multiple comparisons test indicated the differences in spirochete numbers were not statistically significant (ANOVA *p*-value = 0.270, Tukey’s multiple comparisons test of Δd*ksA* fed larvae vs. Δ*dksA* colonized nymphs, *p*-value > 0.999). Cohorts of 5 – 10 *I. scapularis* nymphs infected with either the wild-type, Δ*dksA*, or Δ*dksA* cDksA strains fed upon a total of three mice. Three weeks after the *I. scapularis* bloodmeal, mice were euthanized. The ear, joint, and bladder tissues dissected from euthanized mice were collected and placed into BSK II media. Outgrowth was detected in the tissues collected from mice fed upon by nymphs infected with the wild-type and Δ*dksA* cDksA strains, but not the Δ*dksA* strain (Table 2). Seroconversion was only observed in blood collected from mice infected with wild-type and Δ*dksA* cDksA *B. burgdorferi* strains whose collected tissues showed detectable *B. burgdorferi* outgrowth following culture in BSKII media (Supplemental Fig. 4). These data provide additional evidence that DksA is either required for transmission from *I. scapularis* to the host or, that the Δ*dksA* strain is readily cleared by the host immune system, prior to the generation of a strong antibody response, similar to what is seen for mutants unable to synthesize OspC (Ref).

**Table 2.**
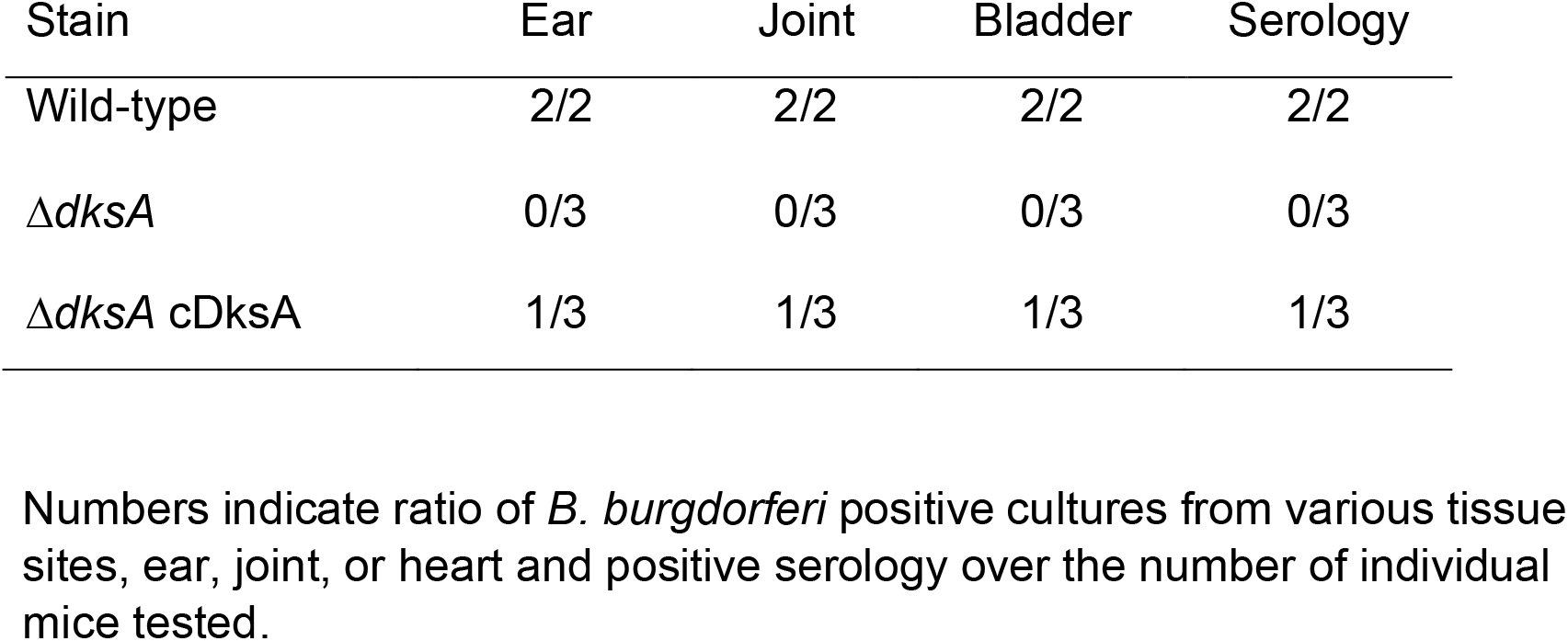
Evaluation of DksA role in *B. burgdorferi* transmission *I. scapularis* nymphs.

### Activation of the *B. burgdorferi* stringent response by changes in pH

The stringent response regulator DksA and Rel_bbu_ synthesized (p)ppGpp control the expression of many genes associated with nutrient uptake and peptide synthesis/protease functions (11, 25). In this study, we assessed the role of DksA and Rel_bbu_ on the expression of RpoS during growth in an acidic medium. To better understand the potential involvement of the stringent response during growth in mildly acidic conditions, we directly analyzed the levels of DksA and (p)ppGpp in *B. burgdorferi*. The relative levels of DksA in wild-type, Δ*relbbu*, Δ*dksA*, Δ*dksA* cDksA strains were analyzed during mid-logarithmic growth in BSKII pH 7.6 and BSKII pH 6.7 + 20 mM acetate by western blot (Fig. 8). DksA protein levels decreased by roughly one-half during growth under the mildly acidic conditions in wild-type cells. DksA expression levels were not significantly different between wild-type, Δ*rel*_bbu_ and the Δ*dksA* cDksA strains. These results suggest the expression of DksA is independent of Rel_bbu_ and chromosomal complementation of the Δ*dksA* strain restored wild-type patterns of DksA expression when strains were grown to mid-logarithmic phase in BSKII pH 7.6 and BSKII pH 6.7 + 20 mM acetate.

**Figure 8.**
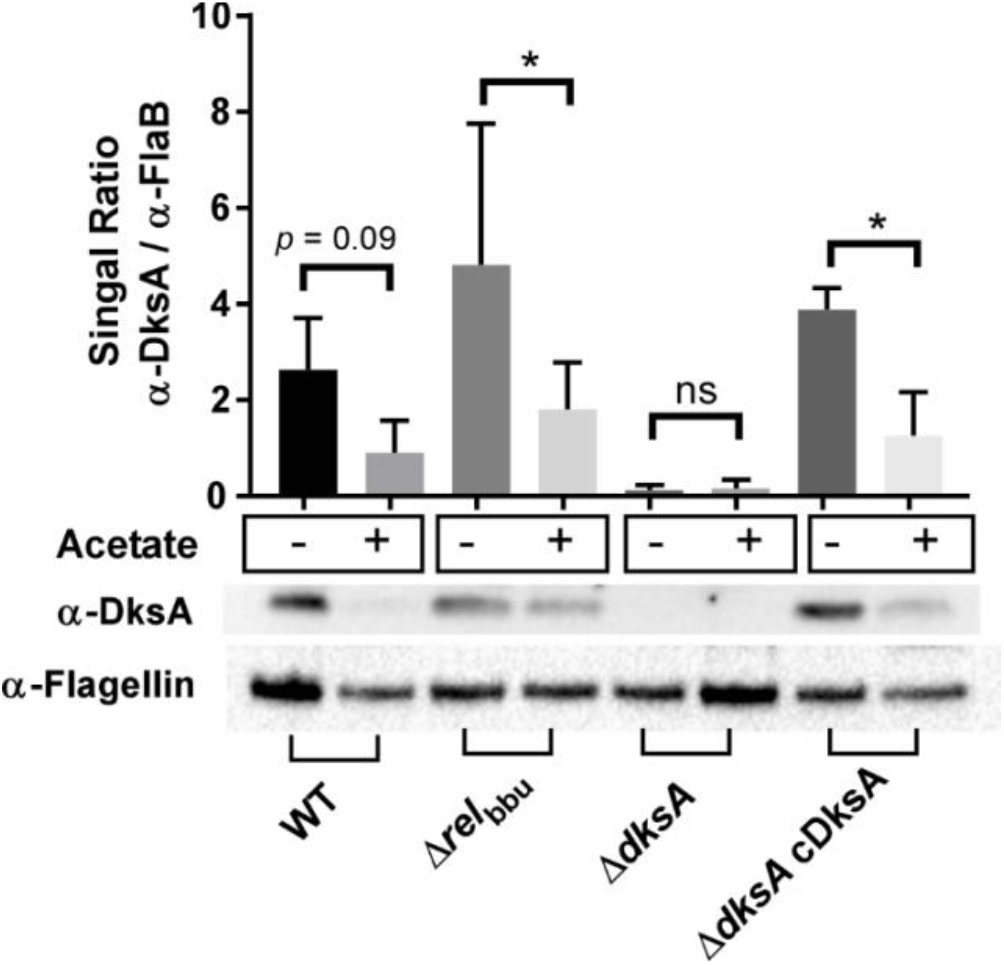
Detection of shifts in DksA protein levels with change in growth media pH and organic acid level at mid-logarithmic growth. Whole cell lysates were collected from 297 wild-type (WT), Δ*rel*_Bbu_, Δ*dksA*, and chromosomally complemented Δ*dksA* (Δ*dksA* cDksA) *B. burgdorferi* strains cultured to 5 × 10^7^ spirochetes *I* mL density in either BSKII-pH 7.6 or BSKII-pH 6.7 with 20 mM acetate. Expression of DksA protein levels **(A)** and OspC protein levels **(B)** were determined by densitometry of chemiluminescence signals and signals were normalized to expression of FlaB. Data are representative of 3 replicate experiments. Asterisk indicate adjusted *p*-values < 0.05 a Sidak multiple comparisons test.

Previously, (p)ppGpp levels were shown to be constitutively elevated in the Δ*dksA* mutant compared to the wild-type strain (11). Since the absence of DksA leads to elevated cellular levels of (p)ppGpp in *B. burgdorferi*, we reasoned the DksA and (p)ppGpp levels may have an inverse relationship. (p)ppGpp is a product of GTP modification and formation of (p)ppGpp can sequester GTP from other cellular processes. To characterize the levels of GTP and (p)ppGpp in *B. burgdorferi* GTP and (p)ppGpp levels were quantified from cell lysates by HPLC. Spirochetes were grown under mildly acidic conditions, wild-type and Δ*dksA* strains were cultured to 1 – 2 × 10^7^ spirochetes · ml^-1^ in either BSK II pH 7.6 or BSK II pH 6.7 with and without 20 mM acetate. Cultures of wild-type and Δ*dksA* strains grown at pH 6.7 had roughly 2-fold lower levels of GTP compared to cultures grown at pH 7.6 (Fig. 9A). Concurrently, both wild-type and Δ*dksA* strains had nearly 10-fold higher levels of (p)ppGpp when cultured at pH 6.7 compared to levels of (p)ppGpp in wild-type spirochetes grown at pH 7.6. GTP levels decrease as (p)ppGpp levels increase in wild-type cells. Additionally, regardless of growth condition, the Δ*dksA* strain had (p)ppGpp levels that were 10-fold higher than wild-type spirochetes grown at pH 7.6, consistent with our previously published observations (11). The ratio of GTP to (p)ppGpp was roughly 70:1 in wild-type cells during growth at pH 7.6, while the GTP to (p)ppGpp ratio was near 2:1 during growth at pH 6.7, meaning a greatly increased molar concentration of (p)ppGpp relative to GTP occurred in cells during growth at pH 6.7. Cellular levels of (p)ppGpp are canonically associated with stationary phase growth and the arrest of cell replication (67). The levels of (p)ppGpp in wild-type *B. burgdorferi* grown to logarithmic or stationary phase in BSKII pH 6.7 or pH 6.7 + 20 mM acetate were also measured by HPLC to determine if increases in (p)ppGpp are associated with growth to stationary phase (Fig. 9C). Results indicate the levels of (p)ppGpp in wild-type cells grown to stationary phase are identical to log phase cells in BSKII pH 6.7 or pH 6.7 + 20 mM acetate. Together these results illustrate *B. burgdorferi* robustly regulates the stringent response in an acidic environment and suggests a relationship between pH and nutrient stress responses in *B. burgdorferi*.

**Figure 9.**
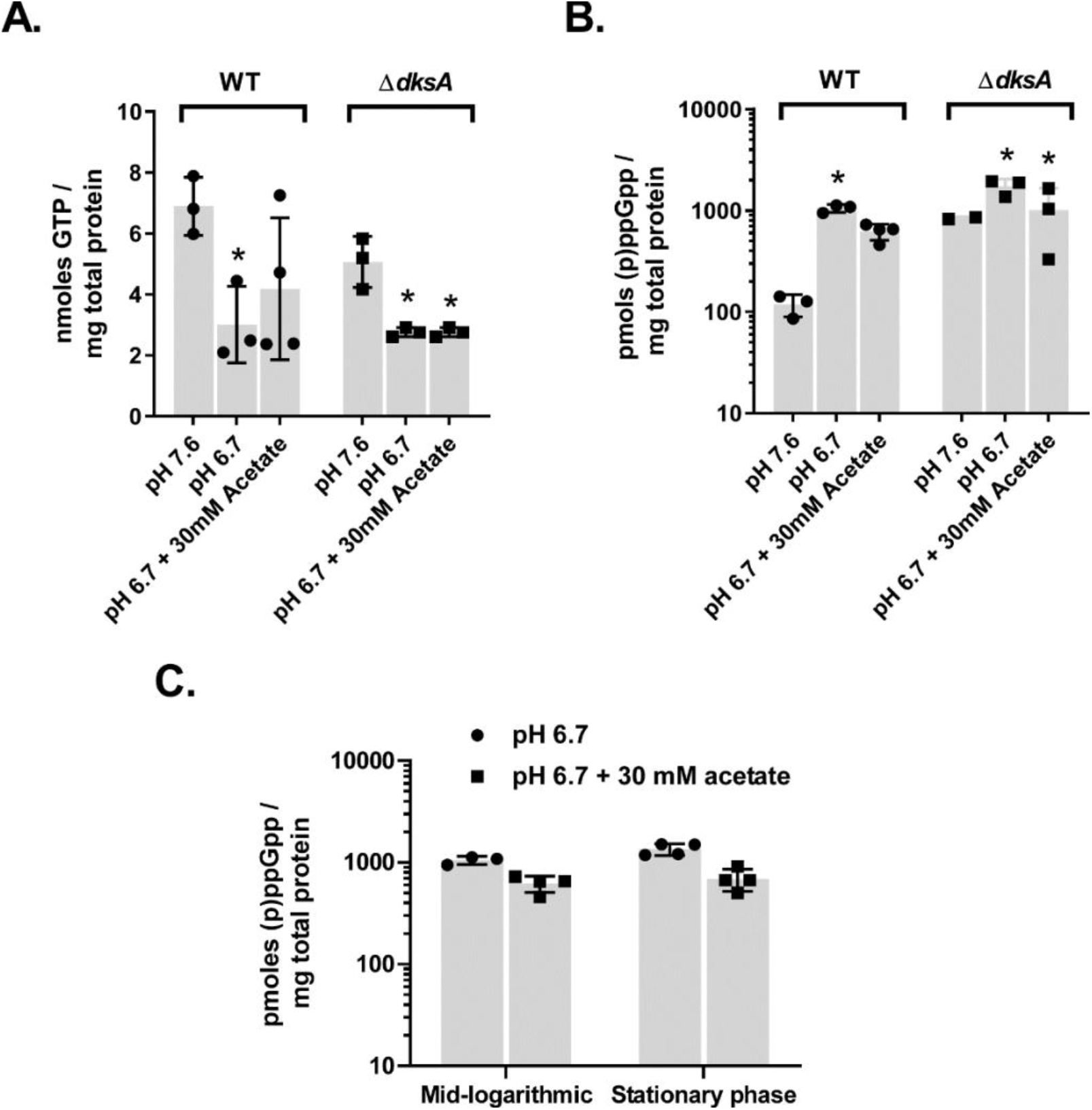
GTP and (p)ppGpp content of *B. burgdorferi* cells in response to culture media pH and organic acid content. Wild-type (WT) and Δ*dksA* strains cultured in BSK II pH 7.6 or in BSK II pH 6.7 with or without 30 mM acetate to 2 – 4 x 10^7^ spirochetes · ml^-1^ were collected to measure (A) GTP and (B) (p)ppGpp content by HPLC. Three replicate experiments were performed. Asterisks indicate *p* < 0.05 for comparison between WT in BSKII 7.6 measurements and all other measurements in a one-way ANOVA with multiple comparisons test on levels of GTP and ppGpp. (C) Wild-type *B. burgdorferi* cultured in BSK II pH 6.7 with or without 30 mM acetate to either mid-logarithmic (2 × 10^7^ spirochetes · ml^-1^) or to stationary (2 × 10^8^ spirochetes · ml^-1^) phase were collected to measure cell density dependent (p)ppGpp content by HPLC. Relative measurement and identity of (p)ppGpp were verified by LC-MS.

## DISCUSSION

In this study, we established the role of DksA in the infectious cycle of *B. burgdorferi* and determined the regulatory relationship between DksA and RpoS, a transcriptional regulator required for murine infection. Mice inoculated with the Δ*dksA B. burgdorferi* strain via intraperitoneal inoculation had no culturable spirochetes 23 days post-infection and failed to develop serum antibodies against *B. burgdorferi*, suggesting the mice cleared the mutant strain prior to the development of an immune response. The clearance of the Δ*dksA B. burgdorferi* strain is likely not due to defects in motility (11). It is more likely due to the inability of this strain to express host-associated lipoproteins regulated by RpoS, such as OspC, which are essential to subvert host immune clearance. These observations are consistent with a recent report by Mason C. et. al. that showed DksA was required for the expression of RpoS-regulated genes and for infection of mice following intradermal inoculation, however, the seroconversion of mice infected with *dksA*-deficient *B. burgdorferi* were not reported in that study (63). While DksA is required for mammalian infection by *B. burgdorferi* following intradermal or intraperitoneal inoculation, it is not essential for colonization of *I. scapularis*. We observed the persistence of Δ*dksA B. burgdorferi* in *I. scapularis* for over 6 weeks following artificial infection of ticks by immersion, suggesting DksA is not required for the long-term survival of *B. burgdorferi* within its tick vector. The absence of DksA does not lead to a defect in stationary-phase-like metabolism when *B. burgdorferi* are nutrient constrained and do not replicate. *I. scapularis* are the natural vectors of *B. burgdorferi* and immunomodulatory effects of *I. scapularis* saliva aid in *B. burgdorferi* transmission (68). Despite the successful colonization and transstadial survival of Δ*dksA B. burgdorferi* in *I. scapularis*, these spirochetes are unable to be successfully transmitted by ticks to naïve mice. These observations demonstrate the essential role of DksA in the natural infectious cycle, extend observations by Mason, et. al. and provide further evidence that DksA plays a critical role in gene regulation during *B. burgdorferi* transmission.

Regulation of RpoS plays a crucial role in infectivity of *B. burgdorferi* (45, 50) with multiple transcriptional regulators (Rrp2, RpoN, BosR, BadR) modulating the transcription of *rpoS* (REFS). In this study, we observed differences in RpoS but did not observe differences in the levels of *rpoS* transcription between our wild-type, Δ*dksA*, and Δ*dksA* cDksA strains. This suggests that the control of RpoS levels by DksA is post-transcriptional and is consistent with the recent report by Mason C. et. al. (63), which suggested the small RNA regulator DsrA and the ClpXP protease system are involved in DksA-dependent RpoS regulation. However, whether ClpXP plays a role controlling RpoS levels is currently unknown. Conversely, our study provides evidence that BBD18 likely contributes to the DksA-dependent regulation of RpoS. BBD18 is a post-transcriptional regulator of RpoS in *B. burgdorferi*, and elevated levels of BBD18 correspond to reduced cellular levels of RpoS. In our previous transcriptomic study and in data provided here, we observed elevated levels of *bbd18* transcripts and BBD18 in Δ*dksA B. burgdorferi* strains, suggesting DksA contributes to the regulation of *bbd18*. Collectively, these data suggest DksA indirectly mediates the regulation of RpoS and RpoS-dependent lipoproteins required for infection by controlling the cellular levels of BBD18. Further studies will be needed to decipher the mechanisms underlying DksA-dependent transcriptional regulation of *bbd18* and other potential post-transcriptional regulators of RpoS.

DksA from *Escherichia coli, Salmonella enterica*, and *Pseudomonas aeruginosa* are all transcriptional repressors (22, 59, 60, 69). Moreover, our previous study indicated DksA represses the transcription of a wide variety of genes in *B. burgdorferi* during *in vitro* culture under nutrient-limiting conditions (11). In this study, we determined the molecular basis of DksA function using our recently established *in vitro* transcription assay. The *in vitro* transcription reactions initiated from seven RpoD-dependent promoters all resulted in lower levels of RNA produced by RNA polymerase with the addition of DksA. The relative quantities of DksA required to observe repression of RNA polymerase activity in the *B. burgdorferi in vitro* transcription assay were similar to those observed in *E. coli* and *S. enterica* RNA-polymerase based assays, where the addition of 2 – 5 μM DksA led to reduced transcription from target promoters (21, 30). The addition of 1 μM *E. coli* DksA to *in vitro* transcription reactions reduced the transcription of *rrnB* P1 by 10-fold, given an acidic assay environment (29). This DksA-dependent change in transcriptional activity is similar to the magnitude of transcriptional changes observed in our *B. burgdorferi in vitro* transcription system with the addition of 1 μM DksA in reactions initiated from *napAp* and *flgBp* dsDNA templates. These observations are consistent with our previously published transcriptomic study comparing wild-type and Δ*dksA B. burgdorferi* strains, indicating DksA constrains global transcription during periods of nutrient limitation.

DksA and (p)ppGpp cooperate in regulating gene expression in other organisms (20, 28). We tested whether both ppGpp and DksA are required for regulating transcription by RNA polymerase from five promoters (*clpCp, glpFp, groLp, napAp*, and *nagAp*) using our *in vitro* transcription system which previous studies suggested were regulated by both Rel_bbu_ and DksA (9, 11, 25). While the impact of DksA on *in vitro* transcription from these promoters was apparent at concentrations as low as 250 nM of DksA, we failed to detect a role for ppGpp in regulating RNA polymerase activity at concentrations as high as 200 μM of ppGpp. While ppGpp did not modulate RNA polymerase activity in this assay, we cannot rule out a role for (p)ppGpp due to the absence of other RNA polymerase binding factors, such as the omega subunit (70, 71). These data indicate DksA is capable of independently regulating RNA polymerase activity in *B. burgdorferi*. Further work is necessary to clarify the role of (p)ppGpp in regulating RNA polymerase function, along with its interaction with DksA.

Present data suggest the role of DksA is divergent from (p)ppGpp in *B. burgdorferi*. Previous studies have shown (p)ppGpp is essential for survival in and transmission from *I. scapularis*, but it is not required for murine infection by needle inoculation. In this study, we showed the Δ*rel*_bbu_ *B. burgdorferi* strain expresses OspC when cultured in BSKII pH 6.8 + 20 mM acetate, while the Δ*dksA* strain does not. Therefore, DksA appears to play a stronger role in controlling the expression of lipoproteins required for mammalian infection than (p)ppGpp. In *E. coli*, (p)ppGpp levels regulate *dksA* transcription (72). However, in *B. burgdorferi*, the regulation of DksA expression under low pH was independent of (p)ppGpp (Fig. 8), indicating the regulatory relationship between (p)ppGpp and DksA in *B. burgdorferi* is not synonymous with *E. coli*. Rather, low levels of DksA correspond closely with elevated cellular levels of (p)ppGpp, and the Δ*dksA B. burgdorferi* strain displayed elevated concentrations of (p)ppGpp regardless of growth condition. The regulatory schemes of DksA and (p)ppGpp do not appear reciprocal, and the roles of DksA and (p)ppGpp as regulators of RpoS are potentially divergent.

DksA and (p)ppGpp are canonically considered regulators of stationary phase growth and have been shown to regulate the transcriptional responses of bacteria to nutritional stress (20). However, in *B. burgdorferi*, DksA and (p)ppGpp are regulated dramatically in response to growth in a slightly acidic environment. While the growth rate of all strains in BSKII pH 7.6 and BSKII pH 6.8 + 20 mM acetate are similar, growth in acidic medium reduced the expression levels of DksA by one-half. Additionally, growth of wild-type cells in BSKII pH 6.8 + 20 mM acetate, led to a drastic shift in intracellular GTP:(p)ppGpp ratios from 50:1 to 2:1. A previous study showed growth of *B. burgdorferi* in BSKII pH 6.8 + 20 mM acetate leads to acidification of the cytosol (5). It is possible this slightly acidic environment is nutritionally stressful for *B. burgdorferi*. For example, maintenance of proton transport during growth in acidic environments is likely linked with nutrient transport in *B. burgdorferi* (REF book chapter). The *B. burgdorferi* genome lacks a recognizable electron transport chain to translocate H^+^ ions from the intracellular space to the periplasm (73). Instead, *B. burgdorferi* is thought to rely on an ATP-dependent proton transport through the V-type ATPase to generate and maintain a proton motive force (5). *B. burgdorferi* also lacks the capacity for amino acid biosynthesis and relies largely on ABC-type oligopeptide transporters for growth (14). ATP is required for both ABC-type transporters and the V-type ATPase, indicating these two processes may compete if ATP pools are limited. Previous transcriptomic studies do not support a role for Rel_bbu_ or DksA in the regulation V-type ATPase genes. However, both Rel_bbu_ and DksA contribute to the expression of ABC-type oligopeptide transporter genes (11, 25). Oligopeptide transporters have also been linked to expression of virulence genes like OspC (74, 75). However, the interplay between the pH environment, ATP levels, oligopeptide transporters, DksA and (p)ppGpp requires further investigation.

Our work establishes DksA as a regulator of *B. burgdorferi* RpoS and uncovers a previously unexplored pH-dependent regulation of (p)ppGpp and DksA. Environmental sensing mechanisms feeding into *B. burgdorferi* virulence gene regulation are poorly understood. In other organisms, Rel and DksA most likely act as sensors of the intracellular environment (16, 29, 60). For example, Rel proteins directly trigger the accumulation of (p)ppGpp in many organisms in response amino acid-starved environments (76), and DksA was recently proposed to be an intracellular sensor of pH (29, 77). Additionally, this study demonstrated the utility of our recently developed *in vitro* transcription assay system to determine the impact of DksA on RNA polymerase-dependent transcription and will facilitate future studies on the mechanisms underlying regulatory activity of DksA and other *B. burgdorferi* transcription factors.

## METHODS

### Recombinant DksA expression

Oligonucleotides encoding an *E. coli* codon-optimized version of *B. burgdorferi dksA* was commercially synthesized (GenScript, Piscataway, NJ, United States). The *dksA* gene was PCR amplified using primers listed in supplemental table 1 and cloned into the NheI site of the pMAL-C5X plasmid expression vector using a Gibson assembly kit (New England Biosciences, Ipswich, MA, United States). Top Shot BL21 (DE3) pLysS Chemically Competent *E. coli* (Invitrogen, Carlsbad, CA, United States) were transformed with the resulting expression vector. For protein expression, overnight *E. coli* cultures passaged 1:200 in Lysogeny broth (LB)-lennox broth containing 2 g · L^-1^ glucose and 100 μg · ml^-1^ ampicillin at 37 °C were grown until the cell density reached OD = 0.5. Cultures were then incubated for an additional 2 hours in the presence of 0.3 mM Isopropyl β-D-1-thiogalactopyranoside. Proteins were purified as described in the pMAL-C5X protein expression system protocols (New England Biosciences, Ipswich, MA, United States). To cleave the MBP-tag from the extracted DksA protein, 2 mM CaCl2 was added to the protein elution fractions. The mixture was incubated overnight with Factor Xa protease at a weight ratio of 1:200 of recombinant protein to protease. To perform cation exchange chromatography, the mixture containing DksA was pushed through a HiTrap SP column (GE Healthcare, Chicago, IL, United States) to bind the DksA protein and subsequently eluted with 50 mM – 1M gradient of sodium chloride. DksA protein was purified to apparent homogeneity when proteins in the elution mixtures where separated by SDS-PAGE and visualized following incubation with Coomassie stain. DksA was stored in 50% glycerol, 1 mM 2-mercaptoethanol, and 25 mM Tris buffer pH 7.5.

### Circular dichroism (CD) spectropolarimetry

For CD analysis, DksA was exchanged into a buffer containing 15 mM sodium phosphate and 100 mM NaCl (pH 7) over a PD-10 Sephadex column (GE Healthcare Bio-Sciences, Pittsburg, PA). DksA protein solutions (12.5 μM, 25 μM, 50 μM, 100 μM and 200 μM) were loaded into a 0.01 cm pathlength quartz cell, and CD spectra were scanned at 0.1 nm increments for wavelengths 180 – 260 nm on the Jasco J-810 polarimeter. CD spectra accumulated over 30 passes at pH 7.0 and 22 °C were averaged and data was visualized in GraphPad Prism software (La Jolla, CA, United States).

### Zn^2+^ release assay

Zn^2+^release from DksA was measured using the metal chelator 4-(2-pyridylazo)resorcinol (PAR) (Sigma-Aldrich, St. Louis, MO, United States) (59). A standard curve consisting of various concentrations of ZnCl2 was incubated with PAR, and absorbance at 500 nm was measured using the Cytation 5 multimode plate reader in order to determine a linear detectable range for the zinc ion release assay. To assay Zn^2+^ release from DksA protein samples, 25 μL of DksA at a 100 μM concentration was exposed to 0 – 1.25 mM H_2_O_2_ in a 96-well assay plate, and free Zn^2+^ was measured following the addition of 25 μL of 2 mM PAR after 0, 30, 60, and 90 minutes of exposure.

### *In vitro* transcription

The *in vitro* transcription reaction was carried out as described previously (58). The reaction contained 0.8 U Ribolock RNase inhibitor (Invitrogen, Carlsbad, CA, United States), 21 nM RNA polymerase, 500 nM RpoD, 2 μCi ATP [α-32P] (PerkinElmer, Waltham, MA, United States), 20 μM ATP, 200 μM GTP, 200 μM CTP, and 200 μM UTP. A preliminary mixture containing reaction buffer, RNA polymerase, and RpoD were incubated for 10 minutes on ice prior to the addition of subsequent components. Transcription was initiated with the addition of double-stranded linear template DNA generated by PCR amplification of genomic DNA segments encoding the transcriptional start site using primers previously described and found on supplemental table 1. The double-stranded DNA templates were added to the *in vitro* transcription reaction to a concentration of 10 nM, and reactions were allowed to proceed for 5 minutes at 37 °C. Transcription was terminated by adjusting the reaction mixture to contain 48% formamide and then incubating the reaction at 65 °C for 5 minute. RNA products were separated by gel electrophoresis in 10% TBE-Urea gels (Invitrogen, Carlsbad, CA, United States) at 180 V for 45 minutes. Gels were placed on a Phosphor Screen (GE Healthcare, Chicago, IL, United States) overnight (16 hours) to detect beta decay from ATP [α-32P] incorporated into the RNA, and the resulting signal was developed into an image using the Typhoon FLA 9500 (GE Healthcare, Chicago, IL, United States). Densitometry measurements were determined with Image Lab 6.0.1. Software (Bio-Rad, Hercules, CA, United States).

### Bacterial strains and culture conditions

A low-passage *B. burgdorferi* 297 strain and respective mutants (Supplemental Table 1) were maintained under microaerobic conditions (5% CO2, 3% O2) in BSK II liquid medium pH 7.6 at 34°C. Cultures started from frozen stocks were passaged twice before performing assays. To assess *B. burgdorferi* responses to environments mimicking the pH of the tick midgut, spirochetes were cultured in pH-adjusted BSK II medium. Mid logarithmic phase (5 × 10^7^ spirochetes · ml^-1^) cultures were passaged by 1:100 dilution directly to pH 6.8 medium with or without various levels of acetate and allowed to reach mid-logarithmic phase.

### Genetic transformation

To assess the phenotype of a Δ*dksA* strain complemented *in cis* with a chromosomal copy of *dksA*, a homologous recombination vector was prepared using a Gibson assembly approach for introducing *dksA* into the 297 Δ*dksA* strain used in a previous study (11, 42, 78). To re-introduce *dksA* to its original genomic location, a 3 kB size fragment surrounding the *dksA* mutagenesis site, including an *aph* gene conferring resistance to kanamycin, was amplified by the primers AG-*bb0168*-F1-5’ and AG-*bb0168*-F2-3’ (Supplemental Table 1). The amplified fragment was cloned into the Zero Blunt TOPO vector (Invitrogen, Carlsbad, CA, United States) and transformed into Top10 *E. coli* with kanamycin selection. The TOPO vector containing the 3 kB fragment was amplified by PCR to generate a 6.5 kB linear fragment using the primers AG-F2R2 and AG-F2F2 to prepare for a three-fragment Gibson assembly using the TOPO-vector as the backbone. A roughly 500 bp long fragment encoding a wild-type copy of *dksA* was amplified by PCR from the wild-type *B. burgdorferi* 297 strain by PCR using the primers *dksA*-gibson-F-F2R2 and *dksA*-gibson-R-*aadA* and a roughly 1 kB fragment encoding a streptomycin resistance cassette (*aadA*) driven by the *flgB* promoter was amplified from the pKFSS1 plasmid by PCR using the *aadA*-gibson-F-*dksA* and *aadA-* gibson-R-*kan* (79). Overlapping oligonucleotides for Gibson cloning (Supplemental Table 1) were encoded on primers and were designed to produce the arrangement *bb0167-dksA-kan-aadA-bb0169* (supplemental Fig. 5). The Gibson assembly was performed per manufacturer instructions with a reaction containing 10-fold molar excess of *dksA* and *aadA* insertion fragments to the 6.5 kB backbone fragment. Top10 *E. coli* was transformed with the Gibson assembly reaction mixture with 50 μg · ml^-1^ spectinomycin selection. Plasmids isolated from Top10 *E. coli* clones were sequenced using primers used during the cloning process. The resulting plasmid, TOPO-297-comp-2B, was electroporated into *B. burgdorferi* 297 Δ*dksA*, and *B. burgdorferi* were plated on semi-solid BSK with 50 μg · ml^-1^ streptomycin for selection. PCR was used to assess the presence of the *dksA* gene, the *aadA* gene, and the plasmid content of transformed *B. burgdorferi* (42, 80). The resulting 297 *dksA* cDksA strain plasmid content was identical to wild-type and Δ*dksA* strains and harbors a wild-type *dksA* allele on the chromosome.

### Mouse model of infection

A mouse model of infection was initiated to test the ability of *B. burgdorferi* strains to cause infection. An established outbred strain of Swiss Webster mice at RML, Rocky Mountain Laboratories (RML) mice, were chosen as the model organism. RML mice are susceptible to *B. burgdorferi* infection, approximating the reservoir host with high practicality due to the well-established animal husbandry protocols. Female 6 – 8-week-old mice were inoculated with *B. burgdorferi* via intraperitoneal injection, as previously described (3, 25). *B. burgdorferi* were enumerated by dark-field microscopy on a Petroff-Hauser Counter prior to inoculation and mice were injected with an inoculum containing 10^5^ *B. burgdorferi*. At 23 days post inoculation, mice were anesthetized, and blood samples were obtained by cardiac puncture during euthanasia. The *B. burgdorferi* infection in mice were evaluated by incubating tissues from the ear, joint, and bladder in BSK II medium containing *Borrelia* antibiotics (2.5 μg / mL Amphotericin, 20 μg / mL Phosphomycin, 50 μg / mL Rifampicin). BSK II cultures were incubated at 34 °C in microaerobic conditions and observed continuously for 4 weeks by dark-field microscopy to detect outgrowth.

### Arthropod transmission model

To test the ability of *B. burgdorferi* strains to transmit from vector to host, *I. scapularis* harboring *B. burgdorferi* were allowed to feed upon mice. To prepare *I. scapularis* colonized with various *B. burgdorferi* strains, two hundred *I. scapularis* larvae were artificially infected with *B. burgdorferi. I. scapularis* larvae and nymphs originating from an uninfected tick colony (National Tick Research and Education Resource, Oklahoma State University) were housed at 22 °C and 95% relative humidity. Egg masses were allowed to mature to larval ticks 4 weeks prior to artificial infection. The culture density of *B. burgdorferi* used for artificial infection was quantified using a Petroff-Hauser Counter under darkfield microscopy, and cultures were diluted to a density of 5 × 10^7^ spirochetes · ml^-1^. Cohorts of *I. scapularis* larval ticks were artificially infected by submersion in 2.0 mL screw cap tubes containing 1.5 mL of *B. burgdorferi* culture in BSK II medium for 2 hours, as previously described (81). Following submersion, ticks were washed twice in phosphate buffered saline pH 8.0 and dried with filter paper.

Artificially infected larval cohorts were allowed to attach to 6 – 8-week-old mice for a bloodmeal. Larval ticks were allowed to molt to obtain *B. burgdorferi* infected *I. scapularis* nymphs. Mice were anesthetized before *I. scapularis* were transferred to the mice with a paint brush. Mice were placed in a wire bottom cage to allow for unattached *I. scapularis* to fall into a live trap. Mice and *I. scapularis* were monitored continuously for 7 days to observe the completion of the bloodmeals by all ticks. Fed larvae were monitored for *B. burgdorferi* colonization and successful molt to nymphs. To quantify *B. burgdorferi* within *I. scapularis* larvae and nymphs, cohorts of ticks were washed in 1.5 mL tubes with solutions of 3% H_2_O_2_, 70% ethanol, and PBS pH 8.0. Washed *I. scapularis* were dissected with forceps on a microscope slide and placed in a tube containing 600 μL BSK II medium. The dissected *I. scapularis* were further disrupted via crushing with a pestle. Volumes of 5 μL, 50 μL, and 500 μL of BSK II medium containing the crushed *I. scapularis* were plated as previously described (25) with semi-solid BSK medium containing 2.5 μg / mL Amphotericin, 20 μg / mL Phosphomycin, and 50 μg / mL Rifampicin.

The transmission of *B. burgdorferi* strains from nymphal *I. scapularis* were tested following the molt of artificially infected larval *I. scapularis* to nymphs. Nymphs were transferred to mice, and the mice were monitored as described above. *B. burgdorferi* infection was determined by the same method described for the needle-inoculation method of infection.

### Transcriptional profiling from cultures

*B. burgdorferi* cultures were pelleted at 4°C, 3,200 x g for 20 minutes. Cell pellets were washed once in HN buffer (10 mM HEPES, 10 mM NaCl, pH 8.0). RNA isolation was performed using RNAzol (Sigma-Aldrich, St. Louis, MO, United States) according to kit instructions. The RNA was quantified by spectrophotometry using a TAKE3 plate in a Cytation 5 multi-mode plate reader (Biotek, Winooski, VT, USA), and RNA quality was assessed by analysis of ribosomal RNA bands visualized following gel electrophoresis by SYBR-safe dye incorporation. cDNA was generated from 500 ng of total RNA from each sample with the High Capacity cDNA Reverse Transcriptase kit (Applied Biosystems, Foster City, CA, United States) following kit instructions. RT-qPCR was performed on the cDNA in the CFX Connect Real-Time PCR Detection System (Bio-Rad, Hercules, CA, United States) using Bullseye EvaGreen Master Mix (MIDSCI, St. Louis, MO, United States) reagents and oligonucleotide primers targeting the gene of interest (Supplemental Table 1). The RT-qPCR data were analyzed using the ΔCq method to indicate copies per 16S rRNA transcript levels.

### Western blotting

Cultures were pelleted at 4 °C, 3,200 x g for 20 minutes. *B. burgdorferi* were washed twice with HN buffer and subsequently lysed in lysis buffer (4% SDS, 0.1M Tris-HCl). SDS-PAGE was performed on the Mini-Tetra System (Bio-Rad, Hercules, CA, United States) using AnykD or 12% polyacrylamide gels and transferred to PVDF membranes using the Transblot Turbo apparatus (Bio-Rad, Hercules, CA, United States). PVDF membranes were blocked for 1-hour in TBST with 5% milk. Commercially available antibodies for OspC (Rockland Immunochemicals, Pottstown, PA), or rabbit polyclonal anti-DksA (11), anti-BBD18 (56), anti-RpoS antibody were incubated with the PVDF membranes overnight at a dilution of 1:2000, 1:2000, 1:500, or 1:500, respectively, in TBST. Antibodies bound to the membrane were detected with the incubation of anti-rabbit HRP-conjugated secondary antibodies for 1 hour followed by five washes in TBST. Images were produced by the ChemiDoc apparatus (Bio-Rad, Hercules, CA, United States) using ECL reagent (LI-COR, Lincoln, NE, United States).

### Quantification of GTP and (p)ppGpp

*Borrelia* cell-free lysates were analyzed for GTP, ppGpp, and pppGpp concentrations by HPLC. Wild-type *B. burgdorferi* 297 and its derivative 297 Δ*dksA* strain were grown to mid logarithmic or stationary phase (1 – 2 × 10^8^ spirochetes · ml^-1^). One hundred ml cultures were centrifuged at 5,000 RPM and washed twice with HN buffer. Cells were lysed by resuspension in distilled water followed by boiling at 99°C for 15 min. Aliquots were removed from each sample for total protein quantification by BCA assay (Thermo Fisher Scientific, Grand Island, NY, United States). After boiling, the suspensions were centrifuged, and the supernatants were extracted twice: first with 2 ml 95% ethanol, second with 1 ml 70% ethanol. The resulting extracts were combined and evaporated under a nitrogen stream at 34°C. Upon evaporation, the samples were resuspended in mobile phase A and filtered through a 0.45 μm nylon membrane syringe filter (GE Healthcare, Chicago, IL, United States). Mobile phase A consisted of 0.15 M triethylammonium acetate (TEAA) pH 5.0, and mobile phase B consisted 0.15M TEAA in acetonitrile. Prior to use, mobile phases were filtered using a 0.2 μm filter with a vacuum followed by ultrasonic degassing. Separations were conducted using a SUPELCOSIL™ LC-18T column, 25 cm x 4.6mm i.d., 5μm particle size (Supelco, Bellefonte, PA, United States). Gradient elution was set to 0 – 4 min, 99% A to 99% A (1 ml/min); 4 – 10 min, 99% A to 85% A (1 ml/min); 10 – 18 min, 85% A to 75% A (0.4 ml/min); 18 – 20 min, 75% A to 25% A (1 ml/min); 20 min – 24 min, 25% A to 97.5% A (1 ml/min); 24 min – 24 min, 97.5% A to 99.0% A (1 ml/min). 20 μl sample injections were used and guanosine triphosphate (Sigma-Aldrich, St. Louis, MO, United States) and guanosine tetraphosphate (Jena Biosciences, Jena, Germany) were used as standards for peak identification and quantification.

### Ethics statement

*In vivo* studies were approved by the Institutional Animal Care and Use Committee of the Rocky Mountain Laboratories (RML). Animal work was conducted adhering to the institution’s guidelines for animal use, and followed the guidelines and basic principles in the United States Public Health Service Policy on Humane Care and Use of Laboratory Animals, and the Guide for the Care and Use of Laboratory Animals by certified staff in an Association for Assessment and Accreditation of Laboratory Animal Care (AAALAC) International accredited facility.

## Author contributions

Conceptualization: William K. Boyle, Crystal L. Richards, Frank C. Gherardini, Travis J. Bourret

Data curation: William K. Boyle, Crystal L. Richards, Daniel P. Dulebohn

Formal analysis: William K. Boyle, Crystal L. Richards

Funding acquisition: Frank C. Gherardini, Travis J. Bourret

Investigation: William K. Boyle, Crystal L. Richards, Daniel P. Dulebohn, Amanda K. Zalud, Jeff A. Shaw

Methodology: William K. Boyle, Crystal L. Richards, Sandor Lovas

Project administration: William K. Boyle, Frank C. Gherardini, Travis J. Bourret Resources: Sándor Lovas, Frank C. Gherardini, Travis J. Bourret

Software: William K. Boyle

Supervision: Crystal L. Richards, Daniel P. Dulebohn, Sandor Lovas, Frank C. Gherardini, Travis J. Bourret

Validation: William K. Boyle, Crystal L. Richards, Amanda K. Zalud, Jeff A. Shaw Visualization: William K. Boyle

Writing – original draft preparation: William K. Boyle

Writing – review & editing: William K. Boyle, Crystal L. Richards, Daniel P. Dulebohn, Amanda K. Zalud, Sándor Lovas, Frank C. Gherardini, Travis J. Bourret

## Acknowledgements

We thank Patricia Rosa for the BBD18 antibody, and Sandy Stewart and Benjamin Schwarz for providing valuable insights and supporting this work.

## References

1. Orloski KA, Hayes EB, Campbell GL, Dennis DT. Surveillance for Lyme disease--United States, 1992–1998. MMWR CDC Surveill Summ. 2000;49(3):1–11.

2. Centers for Disease Control. 2018. Lyme Disease. Available: https://www.cdc.gov/lyme/stats/tables.html. 2018.

3. Bontemps-Gallo S, Lawrence K, Gherardini FC. Two Different Virulence-Related Regulatory Pathways in Borrelia burgdorferi Are Directly Affected by Osmotic Fluxes in the Blood Meal of Feeding Ixodes Ticks. PLoS Pathog. 2016;12(8):e1005791.

4. Bourret TJ, Lawrence KA, Shaw JA, Lin T, Norris SJ, Gherardini FC. The Nucleotide Excision Repair Pathway Protects *Borrelia burgdorferi* from Nitrosative Stress in *Ixodes scapularis* Ticks. Front Microbiol. 2016;7:1397.

5. Dulebohn DP, Richards CL, Su H, Lawrence KA, Gherardini FC. Weak Organic Acids Decrease *Borrelia burgdorferi* Cytoplasmic pH, Eliciting an Acid Stress Response and Impacting RpoN- and RpoS-Dependent Gene Expression. Front Microbiol. 2017;8:1734.

6. Sonenshine DE. Biology of ticks. New York: Oxford University Press; 1991.

7. Zhi H, Xie J, Skare JT. The Classical Complement Pathway Is Required to Control *Borrelia burgdorferi* Levels During Experimental Infection. Front Immunol. 2018;9:959.

8. Stewart PE, Wang X, Bueschel DM, Clifton DR, Grimm D, Tilly K, et al. Delineating the requirement for the *Borrelia burgdorferi* virulence factor OspC in the mammalian host. Infect Immun. 2006;74(6):3547–53.

9. Bugrysheva JV, Pappas CJ, Terekhova DA, Iyer R, Godfrey HP, Schwartz I, et al. Characterization of the RelBbu Regulon in *Borrelia burgdorferi* Reveals Modulation of Glycerol Metabolism by (p)ppGpp. PLoS One. 2015;10(2):e0118063.

10. Drecktrah D, Hall LS, Rescheneder P, Lybecker M, Samuels DS. The Stringent Response-Regulated sRNA Transcriptome of *Borrelia burgdorferi*. Front Cell Infect Microbiol. 2018;8:231.

11. Boyle WK, Groshong AM, Drecktrah D, Boylan JA, Gherardini FC, Blevins JS, et al. DksA Controls the Response of the Lyme Disease Spirochete *Borrelia burgdorferi* to Starvation. J Bacteriol. 2019;201(4).

12. Pappas CJ, Iyer R, Petzke MM, Caimano MJ, Radolf JD, Schwartz I. *Borrelia burgdorferi* requires glycerol for maximum fitness during the tick phase of the enzootic cycle. PLoS Pathog. 2011;7(7):e1002102.

13. Tilly K, Elias AF, Errett J, Fischer E, Iyer R, Schwartz I, et al. Genetics and regulation of chitobiose utilization in *Borrelia burgdorferi*. J Bacteriol. 2001;183(19):5544–53.

14. Groshong AM, Dey A, Bezsonova I, Caimano MJ, Radolf JD. Peptide Uptake Is Essential for Borrelia burgdorferi Viability and Involves Structural and Regulatory Complexity of its Oligopeptide Transporter. mBio. 2017;8(6).

15. Zhu M, Pan Y, Dai X. (p)ppGpp: the magic governor of bacterial growth economy. Curr Genet. 2019;65(5):1121–5.

16. Magnusson LU, Farewell A, Nystrom T. ppGpp: a global regulator in *Escherichia coli*. Trends Microbiol. 2005;13(5):236–42.

17. Traxler MF, Summers SM, Nguyen HT, Zacharia VM, Hightower GA, Smith JT, et al. The global, ppGpp-mediated stringent response to amino acid starvation in *Escherichia coli*. Mol Microbiol. 2008;68(5):1128–48.

18. Sanchez-Vazquez P, Dewey CN, Kitten N, Ross W, Gourse RL. Genome-wide effects on Escherichia coli transcription from ppGpp binding to its two sites on RNA polymerase. Proc Natl Acad Sci U S A. 2019;116(17):8310–9.

19. Holley CL, Zhang X, Fortney KR, Ellinger S, Johnson P, Baker B, et al. DksA and (p)ppGpp have unique and overlapping contributions to *Haemophilus ducreyi* pathogenesis in humans. Infect Immun. 2015;83(8):3281–92.

20. Gourse RL, Chen AY, Gopalkrishnan S, Sanchez-Vazquez P, Myers A, Ross W. Transcriptional Responses to ppGpp and DksA. Annu Rev Microbiol. 2018;72:163–84.

21. Ross W, Sanchez-Vazquez P, Chen AY, Lee JH, Burgos HL, Gourse RL. ppGpp Binding to a Site at the RNAP-DksA Interface Accounts for Its Dramatic Effects on Transcription Initiation during the Stringent Response. Mol Cell. 2016;62(6):811–23.

22. Molodtsov V, Sineva E, Zhang L, Huang X, Cashel M, Ades SE, et al. Allosteric Effector ppGpp Potentiates the Inhibition of Transcript Initiation by DksA. Mol Cell. 2018;69(5):828–39 e5.

23. Lyzen R, Maitra A, Milewska K, Kochanowska-Lyzen M, Hernandez VJ, Szalewska-Palasz A. The dual role of DksA protein in the regulation of *Escherichia coli* pArgX promoter. Nucleic Acids Res. 2016;44(21):10316–25.

24. Paul BJ, Berkmen MB, Gourse RL. DksA potentiates direct activation of amino acid promoters by ppGpp. Proc Natl Acad Sci U S A. 2005;102(22):7823–8.

25. Drecktrah D, Lybecker M, Popitsch N, Rescheneder P, Hall LS, Samuels DS. The *Borrelia burgdorferi* RelA/SpoT Homolog and Stringent Response Regulate Survival in the Tick Vector and Global Gene Expression during Starvation. PLoS Pathog. 2015;11(9):e1005160.

26. Bugrysheva J, Dobrikova EY, Sartakova ML, Caimano MJ, Daniels TJ, Radolf JD, et al. Characterization of the stringent response and rel(Bbu) expression in Borrelia burgdorferi. J Bacteriol. 2003;185(3):957–65.

27. Bugrysheva J, Dobrikova EY, Godfrey HP, Sartakova ML, Cabello FC. Modulation of Borrelia burgdorferi stringent response and gene expression during extracellular growth with tick cells. Infect Immun. 2002;70(6):3061–7.

28. Dalebroux ZD, Svensson SL, Gaynor EC, Swanson MS. ppGpp conjures bacterial virulence. Microbiol Mol Biol Rev. 2010;74(2):171–99.

29. Furman R, Danhart EM, NandyMazumdar M, Yuan C, Foster MP, Artsimovitch I. pH dependence of the stress regulator DksA. PLoS One. 2015;10(3):e0120746.

30. Fitzsimmons LF, Liu L, Kim JS, Jones-Carson J, Vazquez-Torres A. *Salmonella* Reprograms Nucleotide Metabolism in Its Adaptation to Nitrosative Stress. MBio. 2018;9(1).

31. Kvint K, Farewell A, Nystrom T. RpoS-dependent promoters require guanosine tetraphosphate for induction even in the presence of high levels of sigma(s). J Biol Chem. 2000;275(20):14795–8.

32. Merrikh H, Ferrazzoli AE, Lovett ST. Growth phase and (p)ppGpp control of IraD, a regulator of RpoS stability, in Escherichia coli. J Bacteriol. 2009;191(24):7436–46.

33. Lin YH, Chen Y, Smith TC, 2nd, Karna SLR, Seshu J. Short-Chain Fatty Acids Alter Metabolic and Virulence Attributes of *Borrelia burgdorferi*. Infect Immun. 2018;86(9).

34. Carroll JA, Garon CF, Schwan TG. Effects of environmental pH on membrane proteins in *Borrelia burgdorferi*. Infect Immun. 1999;67(7):3181–7.

35. Yang X, Goldberg MS, Popova TG, Schoeler GB, Wikel SK, Hagman KE, et al. Interdependence of environmental factors influencing reciprocal patterns of gene expression in virulent *Borrelia burgdorferi*. Mol Microbiol. 2000;37(6):1470–9.

36. Dunham-Ems SM, Caimano MJ, Eggers CH, Radolf JD. *Borrelia burgdorferi* requires the alternative sigma factor RpoS for dissemination within the vector during tick-to-mammal transmission. PLoS Pathog. 2012;8(2):e1002532.

37. Yang XF, Lybecker MC, Pal U, Alani SM, Blevins J, Revel AT, et al. Analysis of the *ospC* regulatory element controlled by the RpoN-RpoS regulatory pathway in Borrelia burgdorferi. J Bacteriol. 2005;187(14):4822–9.

38. Eggers CH, Caimano MJ, Radolf JD. Analysis of promoter elements involved in the transcriptional initiation of RpoS-dependent *Borrelia burgdorferi* genes. J Bacteriol. 2004;186(21):7390–402.

39. Iyer R, Caimano MJ, Luthra A, Axline D, Jr., Corona A, Iacobas DA, et al. Stage-specific global alterations in the transcriptomes of Lyme disease spirochetes during tick feeding and following mammalian host adaptation. Mol Microbiol. 2015;95(3):509–38.

40. Tilly K, Krum JG, Bestor A, Jewett MW, Grimm D, Bueschel D, et al. *Borrelia burgdorferi* OspC protein required exclusively in a crucial early stage of mammalian infection. Infect Immun. 2006;74(6):3554–64.

41. Dunham-Ems SM, Caimano MJ, Pal U, Wolgemuth CW, Eggers CH, Balic A, et al. Live imaging reveals a biphasic mode of dissemination of *Borrelia burgdorferi* within ticks. J Clin Invest. 2009;119(12):3652–65.

42. Samuels DS, Drecktrah D, Hall LS. Genetic Transformation and Complementation. Methods Mol Biol. 2018;1690:183–200.

43. Stevenson B, Seshu J. Regulation of Gene and Protein Expression in the Lyme Disease Spirochete. Curr Top Microbiol Immunol. 2018;415:83–112.

44. Burtnick MN, Downey JS, Brett PJ, Boylan JA, Frye JG, Hoover TR, et al. Insights into the complex regulation of *rpoS* in *Borrelia burgdorferi*. Mol Microbiol. 2007;65(2):277–93.

45. Smith AH, Blevins JS, Bachlani GN, Yang XF, Norgard MV. Evidence that RpoS (sigmaS) in Borrelia burgdorferi is controlled directly by RpoN (sigma54/sigmaN). J Bacteriol. 2007;189(5):2139–44.

46. Miller CL, Karna SL, Seshu J. *Borrelia* host adaptation Regulator (BadR) regulates rpoS to modulate host adaptation and virulence factors in *Borrelia burgdorferi*. Mol Microbiol. 2013;88(1):105–24.

47. Ouyang Z, Zhou J. BadR (BB0693) controls growth phase-dependent induction of *rpoS* and *bosR* in *Borrelia burgdorferi* via recognizing TAAAATAT motifs. Mol Microbiol. 2015;98(6):1147–67.

48. Boylan JA, Posey JE, Gherardini FC. *Borrelia* oxidative stress response regulator, BosR: a distinctive Zn-dependent transcriptional activator. Proc Natl Acad Sci U S A. 2003; 100(20): 11684–9.

49. Samuels DS. Gene regulation in Borrelia burgdorferi. Annu Rev Microbiol. 2011;65:479–99.

50. Ouyang Z, Blevins JS, Norgard MV. Transcriptional interplay among the regulators Rrp2, RpoN and RpoS in *Borrelia burgdorferi*. Microbiology. 2008;154(Pt 9):2641–58.

51. Boardman BK, He M, Ouyang Z, Xu H, Pang X, Yang XF. Essential role of the response regulator Rrp2 in the infectious cycle of *Borrelia burgdorferi*. Infect Immun. 2008;76(9):3844–53.

52. Groshong AM, Gibbons NE, Yang XF, Blevins JS. Rrp2, a prokaryotic enhancer-like binding protein, is essential for viability of *Borrelia burgdorferi*. J Bacteriol. 2012;194(13):3336–42.

53. Grove AP, Liveris D, Iyer R, Petzke M, Rudman J, Caimano MJ, et al. Two Distinct Mechanisms Govern RpoS-Mediated Repression of Tick-Phase Genes during Mammalian Host Adaptation by *Borrelia burgdorferi*, the Lyme Disease Spirochete. mBio. 2017;8(4).

54. Sarkar A, Hayes BM, Dulebohn DP, Rosa PA. Regulation of the virulence determinant OspC by *bbd18* on linear plasmid lp17 of *Borrelia burgdorferi*. J Bacteriol. 2011;193(19):5365–73.

55. Hayes BM, Dulebohn DP, Sarkar A, Tilly K, Bestor A, Ambroggio X, et al. Regulatory protein BBD18 of the lyme disease spirochete: essential role during tick acquisition? mBio. 2014;5(2):e01017–14.

56. Dulebohn DP, Hayes BM, Rosa PA. Global repression of host-associated genes of the Lyme disease spirochete through post-transcriptional modulation of the alternative sigma factor RpoS. PLoS One. 2014;9(3):e93141.

57. Lybecker MC, Samuels DS. Temperature-induced regulation of RpoS by a small RNA in *Borrelia burgdorferi*. Mol Microbiol. 2007;64(4):1075–89.

58. Boyle WK, Hall LS, Armstrong AA, Dulebohn DP, Samuels DS, Gherardini FC, et al. Establishment of an in vitro RNA polymerase transcription system: a new tool to study transcriptional activation in Borrelia burgdorferi. Sci Rep. 2020;10(1):8246.

59. Henard CA, Tapscott T, Crawford MA, Husain M, Doulias PT, Porwollik S, et al. The 4-cysteine zinc-finger motif of the RNA polymerase regulator DksA serves as a thiol switch for sensing oxidative and nitrosative stress. Mol Microbiol. 2014;91(4):790–804.

60. Crawford MA, Tapscott T, Fitzsimmons LF, Liu L, Reyes AM, Libby SJ, et al. Redox-Active Sensing by Bacterial DksA Transcription Factors Is Determined by Cysteine and Zinc Content. MBio. 2016;7(2):e02161–15.

61. Tapscott T, Kim JS, Crawford MA, Fitzsimmons L, Liu L, Jones-Carson J, et al. Guanosine tetraphosphate relieves the negative regulation of *Salmonella* pathogenicity island-2 gene transcription exerted by the AT-rich *ssrA* discriminator region. Sci Rep. 2018;8(1):9465.

62. Pupov D, Petushkov I, Esyunina D, Murakami KS, Kulbachinskiy A. Region 3.2 of the sigma factor controls the stability of rRNA promoter complexes and potentiates their repression by DksA. Nucleic Acids Res. 2018;46(21):11477–87.

63. Mason C, Thompson C, Ouyang Z. DksA plays an essential role in regulating the virulence of *Borrelia burgdorferi*. Mol Microbiol. 2020.

64. Richards CL, Lawrence KA, Su H, Yang Y, Yang XF, Dulebohn DP, et al. Acetyl-Phosphate Is Not a Global Regulatory Bridge between Virulence and Central Metabolism in Borrelia burgdorferi. PLoS One. 2015;10(12):e0144472.

65. Hubner A, Yang X, Nolen DM, Popova TG, Cabello FC, Norgard MV. Expression of *Borrelia burgdorferi* OspC and DbpA is controlled by a RpoN-RpoS regulatory pathway. Proc Natl Acad Sci U S A. 2001;98(22):12724–9.

66. Jutras BL, Chenail AM, Stevenson B. Changes in bacterial growth rate govern expression of the *Borrelia burgdorferi* OspC and Erp infection-associated surface proteins. J Bacteriol. 2013;195(4):757–64.

67. Zyskind JW, Smith DW. DNA replication, the bacterial cell cycle, and cell growth. Cell. 1992;69(1):5–8.

68. Kasumba IN, Bestor A, Tilly K, Rosa PA. Virulence of the Lyme disease spirochete before and after the tick bloodmeal: a quantitative assessment. Parasit Vectors. 2016;9:129.

69. Petushkov I, Esyunina D, Mekler V, Severinov K, Pupov D, Kulbachinskiy A. Interplay between sigma region 3.2 and secondary channel factors during promoter escape by bacterial RNA polymerase. Biochem J. 2017;474(24):4053–64.

70. Igarashi K, Fujita N, Ishihama A. Promoter selectivity of *Escherichia coli* RNA polymerase: omega factor is responsible for the ppGpp sensitivity. Nucleic Acids Res. 1989;17(21):8755–65.

71. Vrentas CE, Gaal T, Ross W, Ebright RH, Gourse RL. Response of RNA polymerase to ppGpp: requirement for the omega subunit and relief of this requirement by DksA. Genes Dev. 2005;19(19):2378–87.

72. Chandrangsu P, Lemke JJ, Gourse RL. The *dksA* promoter is negatively feedback regulated by DksA and ppGpp. Mol Microbiol. 2011;80(5):1337–48.

73. Fraser CM, Casjens S, Huang WM, Sutton GG, Clayton R, Lathigra R, et al. Genomic sequence of a Lyme disease spirochaete, *Borrelia burgdorferi*. Nature. 1997;390(6660):580–6.

74. Zhou B, Yang Y, Chen T, Lou Y, Yang XF. The oligopeptide ABC transporter OppA4 negatively regulates the virulence factor OspC production of the Lyme disease pathogen. Ticks Tick Borne Dis. 2018;9(5):1343–9.

75. Raju BV, Esteve-Gassent MD, Karna SL, Miller CL, Van Laar TA, Seshu J. Oligopeptide permease A5 modulates vertebrate host-specific adaptation of Borrelia burgdorferi. Infect Immun. 2011;79(8):3407–20.

76. Hauryliuk V, Atkinson GC, Murakami KS, Tenson T, Gerdes K. Recent functional insights into the role of (p)ppGpp in bacterial physiology. Nat Rev Microbiol. 2015;13(5):298–309.

77. Fernandez-Coll L, Potrykus K, Cashel M. Puzzling conformational changes affecting proteins binding to the RNA polymerase. Proc Natl Acad Sci U S A. 2018;115(50):12550–2.

78. Gibson DG, Young L, Chuang RY, Venter JC, Hutchison CA, 3rd, Smith HO. Enzymatic assembly of DNA molecules up to several hundred kilobases. Nat Methods. 2009;6(5):343–5.

79. Frank KL, Bundle SF, Kresge ME, Eggers CH, Samuels DS. *aadA* confers streptomycin resistance in *Borrelia burgdorferi*. J Bacteriol. 2003;185(22):6723–7.

80. Blevins JS, Hagman KE, Norgard MV. Assessment of decorin-binding protein A to the infectivity of *Borrelia burgdorferi* in the murine models of needle and tick infection. BMC Microbiol. 2008;8:82.

81. Policastro PF, Schwan TG. Experimental infection of *Ixodes scapularis* larvae (Acari: Ixodidae) by immersion in low passage cultures of *Borrelia burgdorferi*. J Med Entomol. 2003;40(3):364–70.

